# Psilocybin collapses visual change detection and drives cortical dynamics toward a state of surprise

**DOI:** 10.64898/2026.08.21.745777

**Authors:** Roberto De Filippo, Ryan Gillis, David Wyrick, Mikayla Carlson, Severine Durand, R. Carter Peene, Ahad Bawany, Avalon Amaya, Hannah Belski, Conor Grasso, Warren Han, Jaimie Kenney, Carly Kiselycznyk, Henry Loeffler, Lydia C. Marks, Robyn Naidoo, Benjamin Ouellette, Lucas Suarez, Jackie Swapp, Tye Johnson, Julie Weber, Joshua Wilkes, Peter Groblewski, Allison Williford, Michael Buice, Christof Koch, Irene Rembado, Jérôme A. Lecoq, Torben Ott

## Abstract

Psilocybin profoundly alters visual perception, yet the neuronal mechanisms underlying these effects remain unclear. Here we combined large-scale Neuropixels recordings with cell-type-specific optogenetics in head-fixed mice performing a visual change-detection task. Psilocybin severely impaired task performance without overt motor deficits. In cortex, the drug modestly suppressed activity of layer 5 neurons while preserving representations of image identity. By contrast, psilocybin imposed a 4-Hz oscillation on visually evoked activity that preferentially affected neurons encoding image change rather than image identity. Under psilocybin, expected image repetitions aberrantly recruited change-encoding ensembles and shifted cortical population dynamics towards trajectories normally evoked by genuine stimulus changes. These effects were strongest in somatostatin-expressing (SST) interneurons in visual cortex. The strength of this modulation depended on image structure and was greatest for images with clear, continuous contours, which preferentially recruited change-encoding ensembles. These findings demonstrate that psilocybin drives internally generated cortical surprise signals, providing a circuit mechanism for altered perception in the acute psychedelic state.

---

Psychedelics such as psilocybin profoundly alter visual perception, producing vivid perceptual experiences and shifts in sensory salience[1–3]. Predictive processing frameworks propose that these effects arise from aberrant perceptual inference, suggesting that psychedelics distort the balance between top-down expectations and bottom-up sensory evidence[4, 5] or directly alter the computation of prediction error signals[6]. A common prediction of these models is an altered capacity to distinguish expected from unexpected visual inputs. However, how psychedelics alter cortical circuit dynamics and specific cell types to disrupt this computation remains unknown. To address this question, we used a visual change-detection task in head-fixed mice that directly tests the ability to detect deviations from learned visual expectations[7, 8] on the Allen Institute *OpenScope Brain Observatory*. In this go/no-go task, mice were presented with a continuous series of eight natural images they had been previously trained on, each flashed for 250 ms and separated by a 500 ms gray-screen inter-stimulus interval. After a variable number of image repetitions (“no-change images”), stimulus identity changed (“change image”), and mice reported the deviant image by licking a water spout to obtain reward (Figure 1a). This design dissociates neural coding of stimulus identity from neural coding of image change. We combined large-scale Neuropixels recordings across the mouse visual hierarchy with optogenetic identification of SST interneurons and psilocybin administration. We focused on SST interneurons because of their established roles in lateral subtractive inhibition and proposed involvement in prediction error computations[9–12]. This approach allowed us to test how psilocybin impacts the cell-type-specific representations of expected versus unexpected visual information, a key feature of predictive processing.

**Fig. 1.**
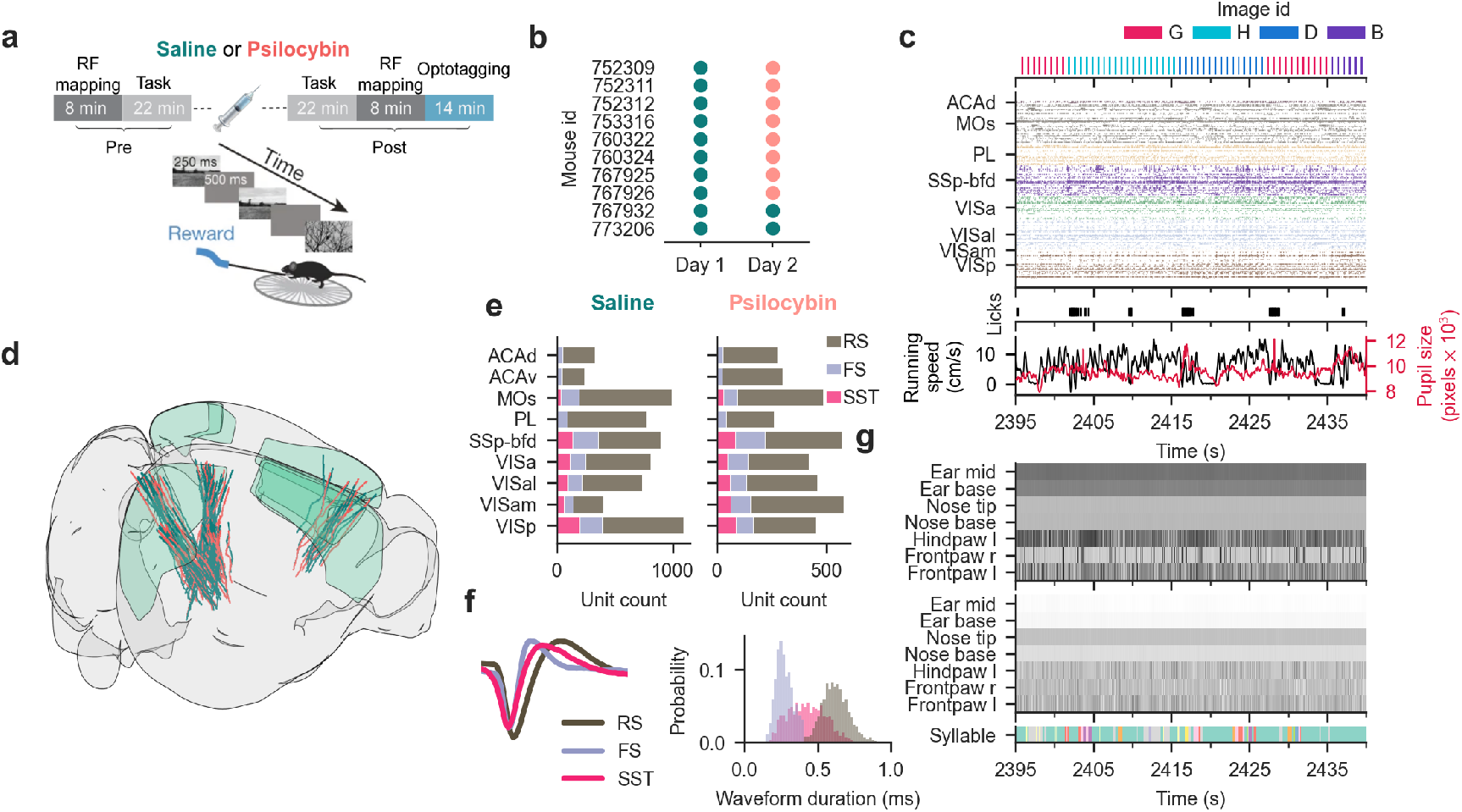
Visual change-detection task and Neuropixels recordings in visual and frontal cortices. (a) Change detection task and experimental design. In brief, head-fixed mice detected image changes in a series of image presentations (250ms with 500ms inter-stimulus interval) by licking a lick spout to collect water reward for correct responses. Receptive field (RF) mapping was conducted before and after task epochs, and opto-tagging of SST interneurons was conducted at the end of each recording session. The behavioral task period was split into a pre-and post-injection epoch, with either saline or psilocybin injections. (b) Experimental design across mice and sessions comprising paired within-animal recording sessions (n=10 mice, n=12 saline and n=8 psilocybin sessions), plus two saline-only controls. Mouse 760322 was excluded from task performance analysis due to chance-level perceptual sensitivity across both sessions. The psilocybin session for mouse 752309 was excluded from task performance and neural analyses due to a data synchronization failure. (c) Forty-five seconds sample of simultaneously recorded behavioral and neural data (image presentations, spiking activity, licking, running speed, and pupil diameter). Anatomical abbreviations: ACAd, Anterior cingulate area dorsal; MOs, Secondary motor area; PL, Prelimbic area; SSp-bfd, Primary somatosensory area, barrel field; VISa, Anterior visual area; VISam, Anteromedial visual area; and VISp, Primary visual area. (d) 3D rendering of all Neuropixels probes, n=71 for saline (teal) and n=48 for psilocybin (salmon) across visual and frontal cortices. (e) A total of 11,012 recorded single-units broken down for cortical area, saline (teal, left) and psilocybin (salmon, right). (f) Left: Average waveform per neuron type. Right: Waveform duration distribution per neuron type, grey for RS, blue for FS and pink for SST. (g) Spatial coordinates (top, medio-lateral axis; middle, ventro–dorsal axis) of seven body-features tracked from the frontal video. On the bottom, quantification of whole-body motor kinematics using MoSeq to identify motor syllables.

## Neuropixels recordings and behavioral tracking during psilocybin administration

To assess the effect of psilocybin on sensory perception, we trained head-fixed mice (n=10) on a well-established operant visual change-detection task implemented as a standardized, high-throughput pipeline at the Allen Institute[7, 13]. Of the 10 mice trained, 9 performed the task successfully on recording days (perceptual sensitivity above chance in pre-injection epochs, one-sample t-test against 0.5). Performance was evaluated across pre-and post-injection epochs over two consecutive days: mice received saline on day 1 and psilocybin (1 mg/kg, intraperitoneal) on day 2, while two additional control animals received saline on both days (Figure 1b). Before each experiment, intrinsic signal imaging was used to map the borders and retinotopy of visual cortices; these maps then guided probe insertions to record from neurons with receptive fields near the center of the visual stimulus. We then recorded single-unit activity using six Neuropixels probes targeting visual and frontal cortical areas (Figure 1e), while simultaneously tracking licking, running speed, pupil diameter, and motor behavior with frontal and lateral cameras (Figure 1c, d, g). After applying quality-control metrics (according to ref.[7]) and a minimum firing rate threshold of 1 spike/s in both behavioral blocks, we obtained an average of 580 *±* 30 and 578 *±* 39 cortical neurons across all probes for psilocybin and saline sessions, respectively, for a total of 11,012 neurons (4,049 psilocybin and 6,963 saline neurons; Figure 1e). Here, we only consider cortical neurons. All mice were of the Sst-IRES-Cre x Ai32 genotype, enabling optotagging of SST interneurons (Figure S1, n = 1,114 optotagged neurons, 62 *±* 5 per session). Fast-spiking (FS) cells were identified based on spike waveform shape (Figure 1f)[14].

## Psilocybin selectively impairs performance in the visual change-detection task but not motor behavior

Psilocybin administration drastically decreased the reward rate (Figure 2a, Figure S2a–b). Both hit and false alarm rates were significantly reduced compared to saline controls (hits: post psilocybin = 0.072 *±* 0.069, n = 7 sessions versus post saline = 0.72 *±* 0.076, n = 12 sessions; false alarms: 0.010 *±* 0.007 versus 0.049 *±* 0.007; mean *±* s.e.m.; two-way repeated-measures ANOVA, condition *×* epoch interaction: P = 0.002 for hits, P <0.001 for false alarms; Figure 2b). While baseline perceptual sensitivity was high and comparable between both groups prior to injection (saline: AUC = 0.93 *±* 0.015; psilocybin: AUC = 0.9 *±* 0.023; P = 0.91, post-hoc Šídák test), psilocybin profoundly reduced discriminability during the post-injection epoch, yielding a significant condition *×* epoch interaction (two-way repeated-measures ANOVA, P = 0.0018; AUC = 0.54 *±* 0.025 for psilocybin versus 0.84 *±* 0.032 for saline; post-hoc comparison: P = 0.0016). Indeed, performance following psilocybin administration was statistically indistinguishable from chance (P = 0.21, one-sample t-test against 0.5). Remarkably, a single animal, out of 7, proved resilient to the drug, maintaining baseline-level task performance following psilocybin administration (denoted by a star ✶ in all plots but included in statistical analyses). Notably, the two saline-saline control animals did not show a performance deficit on the second recording day, suggesting that performance deficits did not arise due to repeated recordings (Figure S2a–b).

**Fig. 2.**
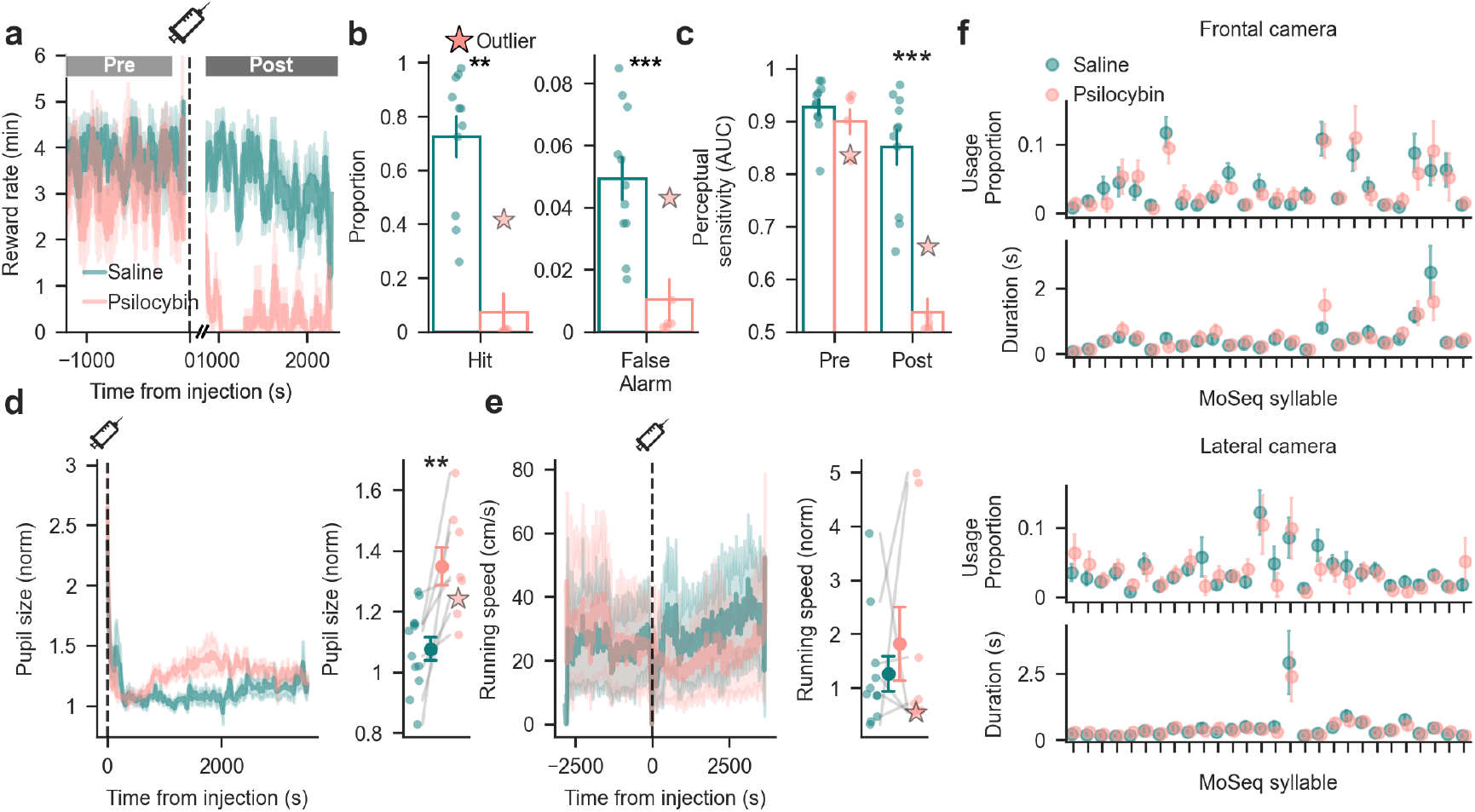
Psilocybin impairs performance in a change detection task. (a) Psilocybin abruptly and significantly impaired change detection performance relative to saline controls. Shading indicates *±*1 s.e.m. (b) Hit and false alarm rate in the post-injection epoch. Performance deficits were driven by reductions in both hit and false alarm rates. Individual points represent single behavioral sessions (n = 11 saline, n = 6 psilocybin). Error bars correspond to *±*1 s.e.m. (c) Perceptual sensitivity (AUC) collapsed to chance levels in the post-injection epoch under psilocybin. Individual points represent single behavioral sessions (n = 11 saline, n = 6 psilocybin). Error bars correspond to *±*1 s.e.m. (d) Psilocybin induced a significant increase in pupil diameter. Within each session, pupil diameter was normalized by dividing it by the mean pupil diameter during the pre-injection baseline period.. Shading indicates *±*1 s.e.m. Individual points represent single behavioral sessions (n = 12 saline, n = 8 psilocybin). (e) Running speed was unaffected by psilocybin administration. Shading indicates *±*1 s.e.m. Individual points represent single behavioral sessions (n = 11 saline, n = 8 psilocybin); running data was unavailable for one session. (f) Motor kinematics (syllable usage proportion and duration) derived from the frontal (top) and lateral (bottom) camera showed no significant differences between conditions (Kruskal-Wallis test, p>0.05 for all syllables). Error bars correspond to *±*1 s.e.m. *P < 0.05, ***P < 0.01, ***P < 0.001.

To assess the effect of psilocybin on the autonomic system, we tracked pupil dynamics. Psilocybin evoked a significant increase in normalized pupil area (psilocybin: 1.349 *±* 0.063, saline: 1.077 *±* 0.039; P = 0.0030, independent t-test; mean *±* s.e.m.; Figure 2d), recapitulating well-established clinical observation of mydriasis in humans[15]. Importantly, the profound disruption of goal-directed behavior occurred without any gross motor deficits. Running speed was unperturbed by the drug (psilocybin: 1.82 *±* 0.68 cm/s, saline: 1.26 *±* 0.32 cm/s; P = 0.90, Mann-Whitney U test; mean *±* s.e.m.; Figure 2e). To capture subtle behavioral phenotypes that might elude macroscopic metrics, we applied an automated motion-sequencing analysis to tracked body parts[16]. This machine-learning approach decomposed multi-feature coordinate trajectories into sub-second, stereotyped behavioral “syllables.” Evaluating the microstructure of motor behavior, we failed to find any discernible differences in syllable composition or usage between groups (Figure 2f, Figure S2c–f). Together, these findings rule out generalized motor impairment, pointing instead to other causes, such as altered visual processing or reduced motivation.

## Psilocybin induces a visual-evoked oscillation strongest in SST interneurons

To investigate the neural basis of the performance deficit, we first asked whether psilocybin altered average firing rates in cortex. To ensure our population analyses strictly isolated the neural correlates of the drug-induced impairment, the single resilient animal was excluded from all following comparisons, leaving n=6 animals undergoing both saline and psilocybin injections. Consistent with previous reports[17–19], psilocybin modestly reduced the firing rates of regular-spiking (RS) neurons relative to saline (P = 0.002, hierarchical bootstrap; Figure S3a–c). Following injection, firing rates increased by 0.09 *±* 0.13 spikes/s under saline but decreased by 1.00 *±* 0.25 spikes/s under psilocybin (mean *±* s.e.m.; P = 0.004, hierarchical bootstrap). By contrast, the firing rates of FS and SST interneurons remained unaltered across all cortical layers (hierarchical bootstrap). To control for potential subtle behavioral confounds not captured by motion-sequencing, we used FaceMap[20] to extract motor principal components and include them as regressors in a generalized linear model (GLM). After accounting for motor variables, residual firing rate differences showed the same pattern, confirming that psilocybin selectively suppressed, cortex-wide, RS neuron activity, while leaving FS and SST firing rates unchanged (on average, Figure S3d–e).

We next asked whether psilocybin altered visual-evoked activity. To isolate visual processing from reward-related confounds that could arise from differences in task performance, we focused our analysis of the effect of psilocybin exclusively on responses to no-change images. Peri-stimulus time histograms (PSTHs) revealed a pronounced 4 Hz rhythmic modulation following psilocybin administration (Figure 3a–b, Figure S4a–b). This oscillation was evident in both spiking activity and local field potentials (Figure S4c). To quantify this rhythmic modulation, we defined an oscillation (Osc) index by fitting a 4-Hz sine wave to the visual-evoked responses (Figure S4d–e). Psilocybin significantly increased the proportion of oscillation-modulated neurons (Osc neurons, P = 0.002, Mann-Whitney U test, Figure 3c). Notably, the single animal that maintained behavioral engagement exhibited minimal induction of these oscillations, with no increase from day 1 to day 2 (Figure 3d, star). Across the population, this robust drug-induced rhythmicity was highly apparent in ΔPSTH heatmaps (Figure 3e, i, m) and in area-averaged ΔPSTHs, with the strongest modulation observed in SST interneurons (Figure 3f, j, n). The modulation was specific to sensory areas (VISp, VISam, VISal, VISa, and SSp-bfd) and was largely absent in frontal regions (Figure 3g, k, o). Due to neuron-type-specific inclusion thresholds, the number of analyzed cortical areas varies across populations (see Methods). Quantification of the Osc index across cortical areas showed that the effect was most pronounced in FS and SST interneurons, and within area VISp (Osc neurons under psilocybin: 35.2% of RS, 45.7% of FS, and 58.9% of SST; Osc neurons under saline: 6.6% of RS, 10.5% of FS, and 9.8% of SST; Figure 3g, k, o). SST interneurons showed the largest group difference in modulation (ΔOsc index=0.55 *±* 0.25, P=0.002, hierarchical bootstrap; Figure 3g, k, o). Notably, while the first two peaks of the modulation coincided with image onset and offset, a third peak became visible during the gray screen period for psilocybin (it was barely visible in saline controls). Together, these results identify a visual-evoked oscillation, strongest in SST interneurons and visual areas, as a signature of psilocybin’s effect on visual processing.

**Fig. 3.**
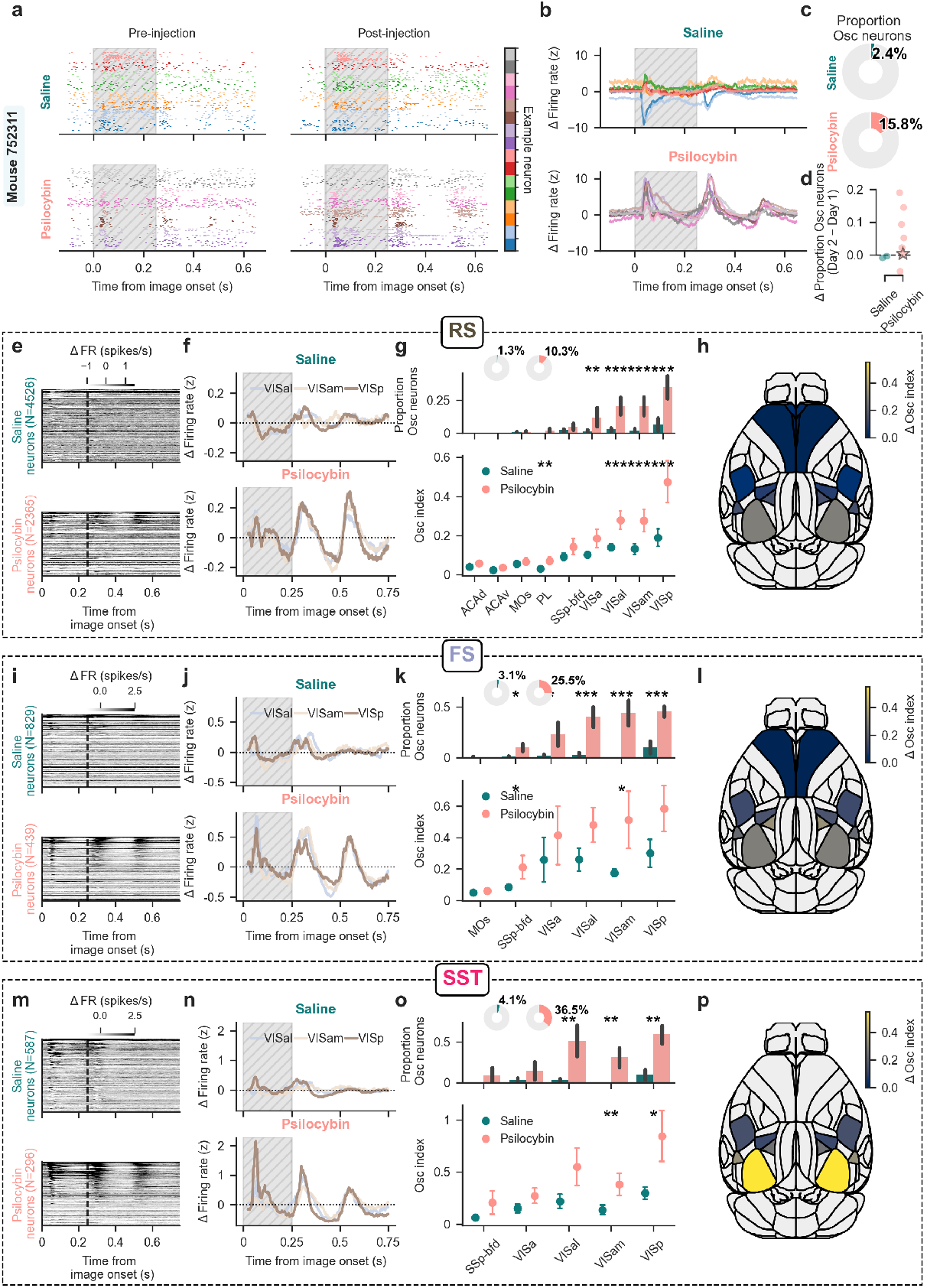
Psilocybin induces an oscillatory modulation strongest in SST interneurons. (a) Example raster plot of 8 representative neurons before (top) and after (bottom) injection of saline (left) or psilocybin (right). The hatched gray area indicates the duration of image presentation. (b) PSTHs of the difference between post-and pre-injection epochs. Each line represents one neuron; neurons from (a) are color-coded accordingly. The hatched gray area indicates the duration of image presentation. (c) Fraction of cortical neurons classified as Osc neurons after injection of saline (top) or psilocybin (bottom). (d) Increase in Osc neurons between experimental sessions. Points represent individual animals. (e) Heatmaps of the change in firing rate (Δ Firing rate = post -pre) for RS neurons, sorted by the Osc index. Data are shown for saline (top) and psilocybin (bottom) groups, aligned to image onset. The dashed line indicates the end of image presentation. (f) Average ΔFiring rate traces for VISp, VISam, and VISal, the areas exhibiting the strongest modulation following psilocybin. The hatched gray area indicates the duration of image presentation. (g) Top, fraction of Osc neurons in each cortical area. P: ACAd, 1.0; ACAv, 1.0; MOs, 1.0; PL, 0.65; SSp-bfd, 0.24; VISa, 0.010; VISal, 0.001; VISam, 0.001; VISp, 0.001, hierarchical bootstrap; pie charts indicate the total percentage of Osc neurons per group. Bottom, distribution of mean Osc index across areas. P: ACAd, 0.090; ACAv, 0.10; MOs, 0.29; PL, 0.002; SSp-bfd, 0.17; VISa, 0.068; VISal, 0.001; VISam, 0.001; VISp, 0.001, hierarchical bootstrap. Error bars correspond to *±*1 s.e.m. (h) Brain map showing the difference in mean Osc index between psilocybin and saline sessions. (i)-(l) Same as E-H for FS neurons. In (k), top P: MOs, 0.55; SSp-bfd, 0.030; VISa, 0.56; VISal, 0.10; VISam, 0.012; VISp, 0.068, hierarchical bootstrap. Bottom, P: MOs, 1.0; SSp-bfd, 0.032; VISa, 0.046; VISal, 0.001; VISam, 0.001; VISp, 0.001, hierarchical bootstrap. (m)-(p) Same as E-H for SST neurons. In (o), top P: SSp-bfd, 0.20; VISa, 0.17; VISal, 0.10; VISam, 0.004; VISp, 0.016, hierarchical bootstrap. Bottom, P: SSp-bfd, 0.64; VISa, 0.25; VISal, 0.006; VISam, 0.002; VISp, 0.002, hierarchical bootstrap. \**P* < 0.05, \*\*\**P* < 0.01, \*\*\**P* < 0.001.

## Psilocybin selectively recruits change-encoding neurons during image repetitions

To isolate the neural signals most relevant to the task, we used mutual information during the pre-injection baseline to identify distinct subpopulations of neurons encoding either image identity (MI_id_) or image change (MI_change_). Because both task-relevant visual representations and the drug-induced oscillatory modulation (Figure S5a–b) were predominantly localized within visual cortical areas (VISa, VISal, VISam, and VISp), we focused all subsequent analyses on these regions. To minimize the influence of licking on the encoding of change (MI_change_), we restricted this analysis to the first 100 ms following image onset. Within the four visual areas we recorded from, RS neurons strongly encoded image identity (Figure 4a, b), whereas image change encoding was significantly enriched in interneurons (Figure 4d, e), consistent with previous reports[21]. Because both MI_id_ and MI_change_ distributions were highly skewed, we defined functional ensembles (id-and change-encoding) using the top 20th percentile of each population. This approach isolated the minority of highly tuned neurons occupying the long tail of each distribution (Figure S5c). The degree of overlap between id-and change-encoding populations depended on neuron type. Specifically, of the neurons classified as highly-tuned change-encoding, only 13.8% of FS and 15.1% of SST interneurons co-represented image identity, whereas RS cells showed a 34.8% overlap (Figure S5d). This change-related signaling was independent of licking behavior, as mean firing rates during no-change trials were indistinguishable between false alarms and correct rejections (Figure S5e–f).

**Fig. 4.**
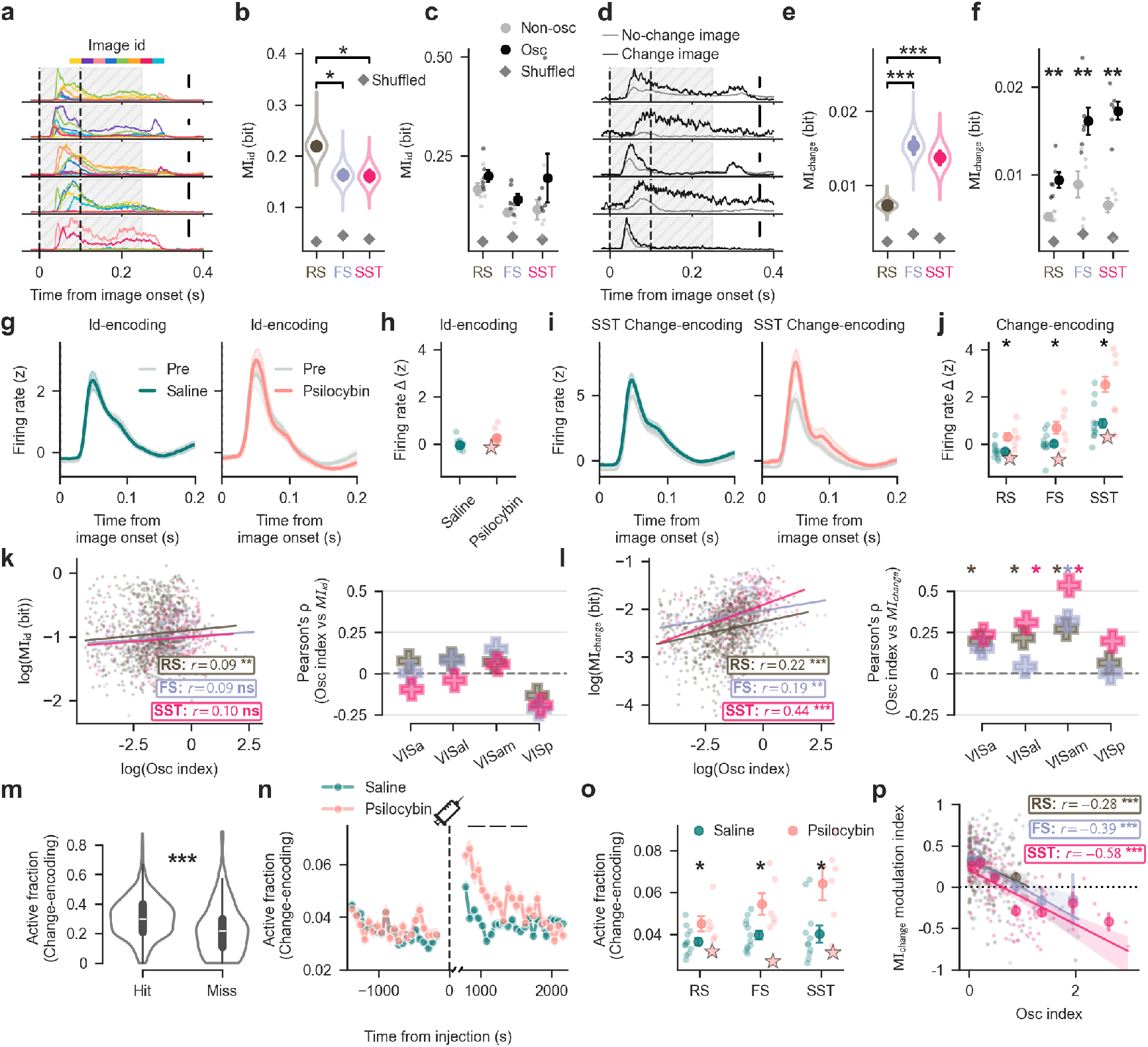
Psilocybin preferentially modulates change-encoding neurons. (a) Five representative single-units encoding image id from one session. Vertical dashed lines indicate the 0–100 ms window used to compute MI. The hatched gray area indicates the duration of image presentation. Scale bar 40 spikes/s. (b) Distribution of MI_id_ values across neuron types (right), with 3 bits necessary to encode image id. RS neurons exhibit an enrichment in MI_id_ (RS = 0.220 *±* 0.004 bits versus FS = 0.163 *±* 0.005 versus SST = 0.162 *±* 0.005; mean *±* s.e.m.; P = 0.048 for RS vs FS, P = 0.042 for RS vs SST; hierarchical bootstrap nested by session and brain area). (c) No significant differences in MI_id_ in Osc neurons (Mann-Whitney U test, P=RS: 0.13, FS: 0.24, SST: 0.31). Error bars correspond to *±*1 s.e.m. (d) Five representative single-units encoding image change from the same session as in A. (e) Distribution of MI_change_ values across SST-inhibition based categories (left) and neuron types (right), with 1 bit necessary to encode an image change. FS and SST neurons exhibit an enrichment in MI_change_ (RS = 0.0074 *±* 0.0001 bits versus FS = 0.0154 *±* 0.0006 versus SST = 0.0138 *±* 0.0005; mean *±* s.e.m.; P <0.001 for RS vs FS, P <0.001 for RS vs SST, hierarchical bootstrap nested by session and brain area). (f) MI_change_ is enriched in psilocybin modulated cells across neuron types (Mann-Whitney U test, P=RS: 0.002, FS: 0.004, SST: 0.002). Psilocybin-modulated cells are defined as the ones with an Osc index in the top 20% percentile. Error bars correspond to *±*1 s.e.m. (g) Firing rate of image id-encoding neurons during no-change trials pre-injection (gray) and post-injection (left: saline, teal; right: psilocybin, salmon). (h) Change in firing rate induced by saline or psilocybin injection, quantified in a 10 ms window centered at the peak response. (i) Firing rate of SST change-encoding neurons during no-change trials pre-injection (gray) and post-injection (left: saline, teal; right: psilocybin, salmon). (j) Change in firing rate induced by saline or psilocybin injection across neuron types, quantified in a 10 ms window centered at the peak response. All neuron types exhibited a significant difference. (k) Correlation between Osc index and MI_id_. Left: Pearson correlation pooled across areas. Only RS neurons showed a significant, weak positive correlation (RS: r = 0.09, P = 0.007). Right: Within-area correlation. No region-cell type pair survived FDR correction (P >0.05, two-sided permutation test). (l) Correlation between Osc index and MI_change_. Left: Pearson correlation pooled across areas. All cell types showed significant positive correlations, strongest in SST neurons (RS: r = 0.22; FS: r = 0.18; SST: r = 0.44; all P <0.005). Right: Within-area correlation. Positive correlations survived FDR correction (P <0.05, two-sided permutation test) for all cell types in VISam, RS in VISa, and RS/SST in VISal. (m) The active fraction of change-encoding neurons during change trials was significantly higher for hits than for misses during pre-injection. (n) Time course of the active fraction of change-encoding neurons during no-change images before and after injection. Psilocybin temporarily increases the active fraction. Grey rectangles represent significance calculated on 5-minute windows (window 1: P=0.022; window 2: P=0.035; window 3: P=0.043; independent t-test). (o) Average active fraction in the 30 min post-injection, shown by neuron type. The star (*✶*) denotes the behavioral outlier. Each dot represents one session. (p) Pearson correlation between the MI_change_ modulation index, quantifying discriminability across epochs of no-change responses compared to pre-injection change responses, and Osc index across change-encoding neurons in psilocybin sessions, shown separately for each neuron type. *P < 0.05, ***P < 0.01, ***P < 0.001.

We next investigated whether Osc neurons were enriched for the encoding of either image identity or image change. Compared to the remaining population, all Osc neurons carried significantly higher change information MI_change_ across all cell types (Figure 4f; RS: P = 0.002; FS: P = 0.004; SST: P = 0.002; Mann-Whitney U test), whereas image id information MI_id_ was indistinguishable between groups (Figure 4c). This difference was consistent across individual brain areas, and especially strong in layer 2/3 (Figure S6).

Examination of PSTHs aligned to no-change trials revealed that psilocybin selectively enhanced the visual onset response of change-encoding neurons, an effect most pronounced within the SST population (Figure 4i, j, Figure S7a–c). Quantifying the injection-induced change in z-scored firing rate within a 10 ms window centered at the peak response confirmed significant elevations under psilocybin relative to saline across all functional cell types (RS: saline = -0.30 *±* 0.07, psilocybin = 0.30 *±* 0.15, P = 0.0010; FS: saline = 0.02 *±* 0.09, psilocybin = 0.69 *±* 0.25, P = 0.014; SST: saline = 0.88 *±* 0.18, psilocybin = 2.52 *±* 0.32, P = 0.0060; bootstrap test). Crucially, this drug-induced enhancement did not apply to image id-encoding neurons, which showed no significant differences in their onset responses to repeated images between the saline and psilocybin conditions (hierarchical bootstrap, P >0.05, Figure 4g, h).

Surprisingly, these task-related signals were related to psilocybin-induced oscillations: a higher degree of image change encoding predicted stronger psilocybin-induced oscillations, with signifi-cantly weaker correlation observed for identity encoding (Figure 4k, l). This continuous scaling was prominent in SST neurons (Pearson’s r = 0.44) and peaked within VISam (Pearson’s r = 0.61), a visual area at the apex of the cortical hierarchy[7, 22]. Significant correlations also emerged in RS neurons within VISal and VISam, and FS neurons in VISam (Figure 4l). Together, these results suggest that an individual neuron’s capacity to encode visual change is related to its susceptibility to oscillatory modulation by psilocybin.

To assess whether the psilocybin-induced increase in image change-encoding neurons is relevant for visual detection, we investigated single-trial dynamics of neural ensembles. We calculated the *active fraction* of id-encoding and change-encoding neurons, respectively, defined as the proportion of neurons whose response to a given image (0 to 100 ms) exceeded their pre-injection average no-change response (see Methods). To assess the behavioral relevance of this metric, we compared the active fraction of change-encoding neurons between hit and miss trials before injection (pre-injection psilocybin and saline were aggregated into a single epoch). We reasoned that hit trials, representing successful detections, should exhibit greater change-encoding activity. In contrast, misses are inherently heterogeneous, potentially arising from either perceptual failures (lack of detection) or motivational lapses (detection without response). Since motivation-driven misses would presumably still engage change-encoding circuits, any hit-miss difference we observe likely underestimates the true magnitude of the perceptual signal. Despite this conservative bias, hit trials exhibited a significantly higher active fraction than misses (Figure 4m; hit: 0.33 *±* 0.03; miss: 0.23 *±* 0.02; P = 0.001, Mann-Whitney U test), confirming that this neural index tracks behavioral performance. Notably, even on hit trials, only a portion of change-encoding neurons were recruited, and the magnitude of this recruitment varied significantly across individual images (P <0.001, Friedman test). This sparse recruitment may reflect the underlying structure of the neural manifold, whereby individual neurons encode only specific image transitions (e.g., Image *A → B* but not Image *C → B*), although the present dataset does not sample sufficient transitions to test this directly. Following psilocybin administration, the active fraction during repeated, no-change images rose, on average, to 56% for approximately 30 min (Figure 4n). This increase was observed across all three neuron types, but was strongest in SST interneurons (Figure 4o; RS: P = 0.041; FS: P = 0.035; SST: P = 0.02). Notably, this aberrant engagement was restricted to change-encoding circuitry; image id-encoding ensembles, excluding neurons with dual selectivity for both identity and change, exhibited no such increase (Figure S8c–e). Indeed, image id encoding MI_id_ remained stable in id-encoding cells before and after psilocybin administration (Figure S8a–b), consistent with reported preservation of visual selectivity under psilocybin[19]. Receptive field mapping indeed confirmed that spatial tuning properties were unaltered post-injection (Figure S9). In contrast, change-encoding neurons, particularly the SST population, showed a strong negative correlation between the magnitude of oscillatory modulation and the discriminability between pre-injection change responses and post-injection no-change responses (Figure 4p; RS: r = -0.25, P <0.001; FS: r = -0.36, P <0.001; SST: r = -0.59, P <0.001). This reveals that elevated oscillatory modulation tracks a loss of discriminative power, impairing a cell’s ability to distinguish expected from unexpected visual inputs. Strikingly, under psilocybin, individual change-encoding neurons exhibited stronger responses to a repeated image than they did to a genuine stimulus change prior to injection, a response reversal observed most prominently within the SST population (Figure S7d–f). In these example neurons, however, post-injection change stimuli evoked even higher firing rates, preserving their relative contrast. Within the change-encoding SST population, the degree of oscillatory modulation significantly correlated with an increase in image identity encoding across all cell types (RS: r = 0.16, P = 0.017; FS: r = 0.21, P = 0.016; SST: r = 0.35, P <0.001). Because this psilocybin-induced modulation falls within the 100 ms window critical for decision-making[13], it likely alters visual processing essential for task execution.

To determine whether these neural activity changes were sufficient to disrupt task-variable decoding, we implemented a cross-epoch decoding analysis targeting both image id and image change. To account for session to session variability, we evaluated two decoders: an intra-epoch decoder using 5-fold cross-validation within the pre-injection baseline, and a cross-epoch decoder trained on pre-injection activity and tested on post-injection activity. Importantly, psilocybin-induced modulation significantly degraded cross-epoch decoding accuracy specifically for distinguishing change versus no-change responses, whereas image id decoding performance was not significantly altered (Figure S10b–f). Because statistical decoders optimize over high-dimensional population variance while completely ignoring the biological constraints that restrict downstream information flow[23], this observed reduction likely underestimates the functional disruption. For the change decoder, significant performance reductions were observed across all functional neuron types, and were selectively localized within area VISam and cortical layer 5 (Figure S10c–g).

## Psilocybin shifts population dynamics toward sensory surprise

We next asked whether the altered activity of change-encoding neurons was accompanied by a reorganization of visual cortical population dynamics, bridging our findings across scales. We used principal component analysis (PCA) to track the temporal evolution of population dynamics in a low-dimensional state space following image onset. We hypothesized that the aberrant activation of change-encoding neurons contributes to shifting the entire neuronal population’s no-change responses toward states normally evoked by genuine image changes. To test this, we focused on early image–onset responses (0–100 ms) and applied PCA to pre–injection data, defining a fixed, low-dimensional subspace that captures the dominant response structure of visual cortex to both change and no–change images. Post–injection no-change responses were then projected into the same PCA space, allowing us to track how psilocybin altered population trajectories. Neural trajectories for change and no-change images diverged across all cell types, as expected[7] (Figure S11). To compare the effects of the injection with genuine image changes on these trajectories, we defined two state-space vectors: the change shift (the difference between pre-injection change and no-change responses), capturing the population activity induced by a true image change, and the injection shift (the difference between post-and pre-injection no-change responses), measuring the drug-induced modulation. While saline control vectors remained near the origin, indicating that the activity remained largely unchanged following saline injections, psilocybin induced a distinct population shift (Figure 5a). In SST interneurons, this trajectory qualitatively mirrored the path of genuine image changes. To determine whether this displacement aligned with the trajectory of a true stimulus transition, we computed in each session the scalar projection of the injection shift vector onto its corresponding change shift vector to yield a change-alignment index. For both RS and SST neurons, psilocybin significantly increased this index relative to saline controls (RS: P = 0.0064; SST: P = 0.044) and temporally shuffled baselines (RS: P = 0.0088; SST: P = 0.035; permutation tests, Figure 5b). Furthermore, in the psilocybin condition, change-encoding SST interneurons exhibited, up to 50 ms after image onset, dynamics that matched the precise temporal profile of true image change responses (Figure 5c). Thus, psilocybin transiently redirects cortical population trajectories by driving no-change responses along a pathway that recapitulates genuine visual transitions, an effect most pronounced within SST interneuron populations.

**Fig. 5.**
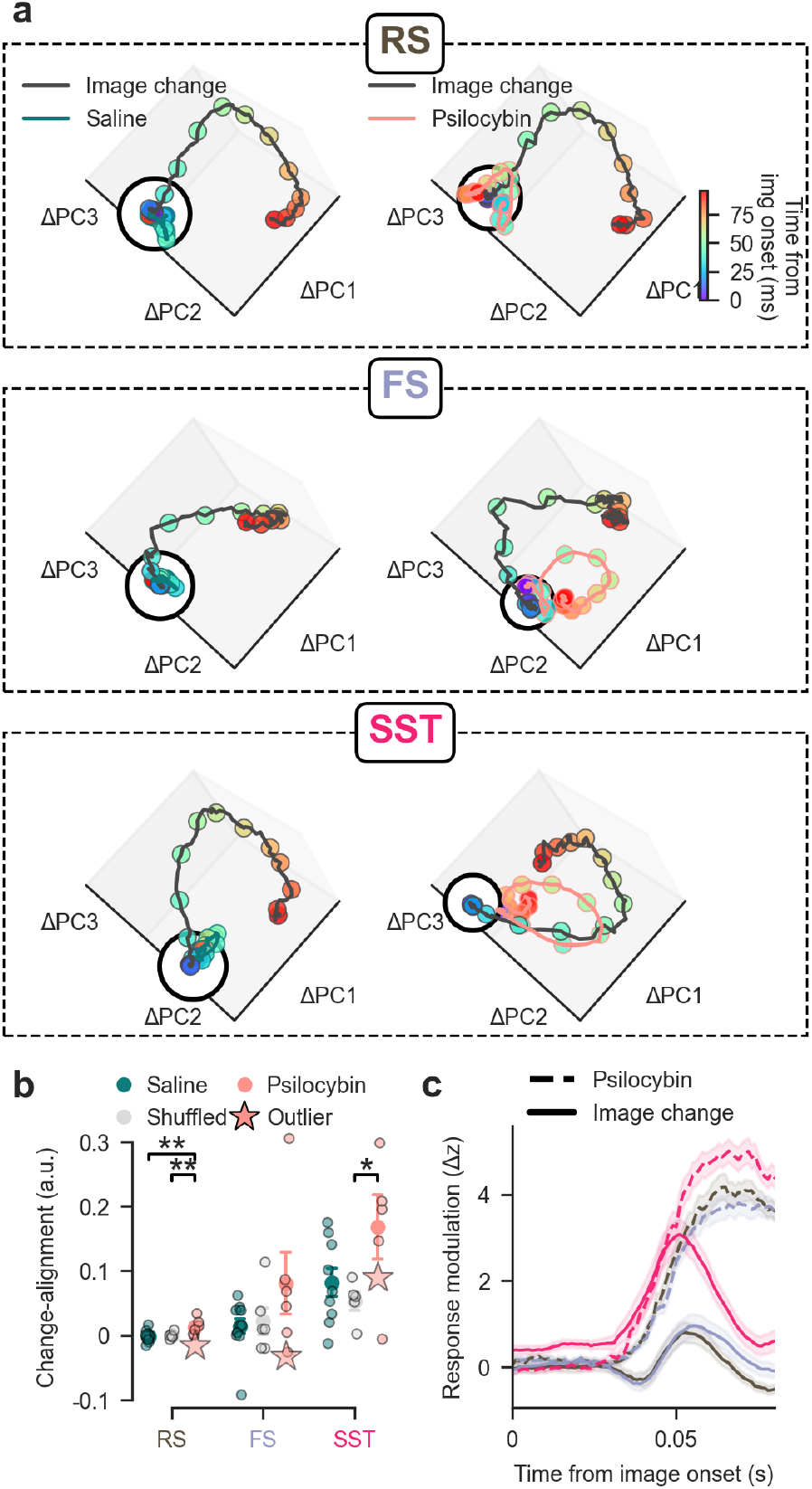
Psilocybin shifts population dynamics in visual cortex towards change-like states. (a) Population trajectory shifts induced by image changes (black line) or injections of saline (teal, left) or psilocybin (salmon, right), projected into a common neural state space. Shifts are calculated relative to baseline activity during pre-injection no-change trials. Dots are color-coded by post-stimulus time (0-100 ms). Rows correspond to neuron types; the black circle denotes the origin. (b) Scalar projection of the injection-effect vector onto the change-effect vector across sessions (see text for details). Psilocybin increases trajectory alignment with genuine image changes in RS and SST neurons, whereas saline and shuffled controls do not. The star (*✶*) denotes the behavioral outlier. Each dot represents one session. Error bars correspond to *±*1 s.e.m. *P < 0.05, ***P < 0.01, ***P < 0.001. (c) Response kinetics of the pre-injection image-change response (Image change, solid lines) and the post-psilocybin no-change response (Psilocybin, dashed lines) in change-encoding neurons, aligned to image onset. Pre-injection no-change responses were subtracted from both conditions to isolate baseline-independent dynamics. Cell types are color-coded as in B. Shading indicates *±*1 s.e.m.

## Stimulus features shape psilocybin-induced modulation

Our preceding analyses treated neural responses to the image set collectively by averaging across all eight images. But does psilocybin reconfigure cortical networks uniformly, or do specific stimulus features dictate this modulation? This question is especially salient given that change-encoding ensembles are differentially recruited across individual images. Indeed, we found that the magnitude of psilocybin-induced oscillatory modulation varied substantially across the image set (Figure 6a). Visual inspection suggested that this variability was related to structural image properties, particularly contour length and density. To quantify this relationship, we defined a Contour Fragmentation Index (CFI) as the ratio of the number of discrete contours to their average length. Under this metric, smooth and continuous contours yield CFI values near zero, whereas highly fragmented contours produce correspondingly large CFI values. CFI accounted for, on average, 56.1% of the within-session variance in Osc index across images during psilocybin sessions (mean r = -0.78, P=0.0017; Figure 6b, c). A similar but weaker relationship was observed during saline sessions, in which CFI accounted for an average of 31.7% of the within-session variance (mean r = -0.56, P<0.001), despite substantially smaller Osc-index values.

**Fig. 6.**
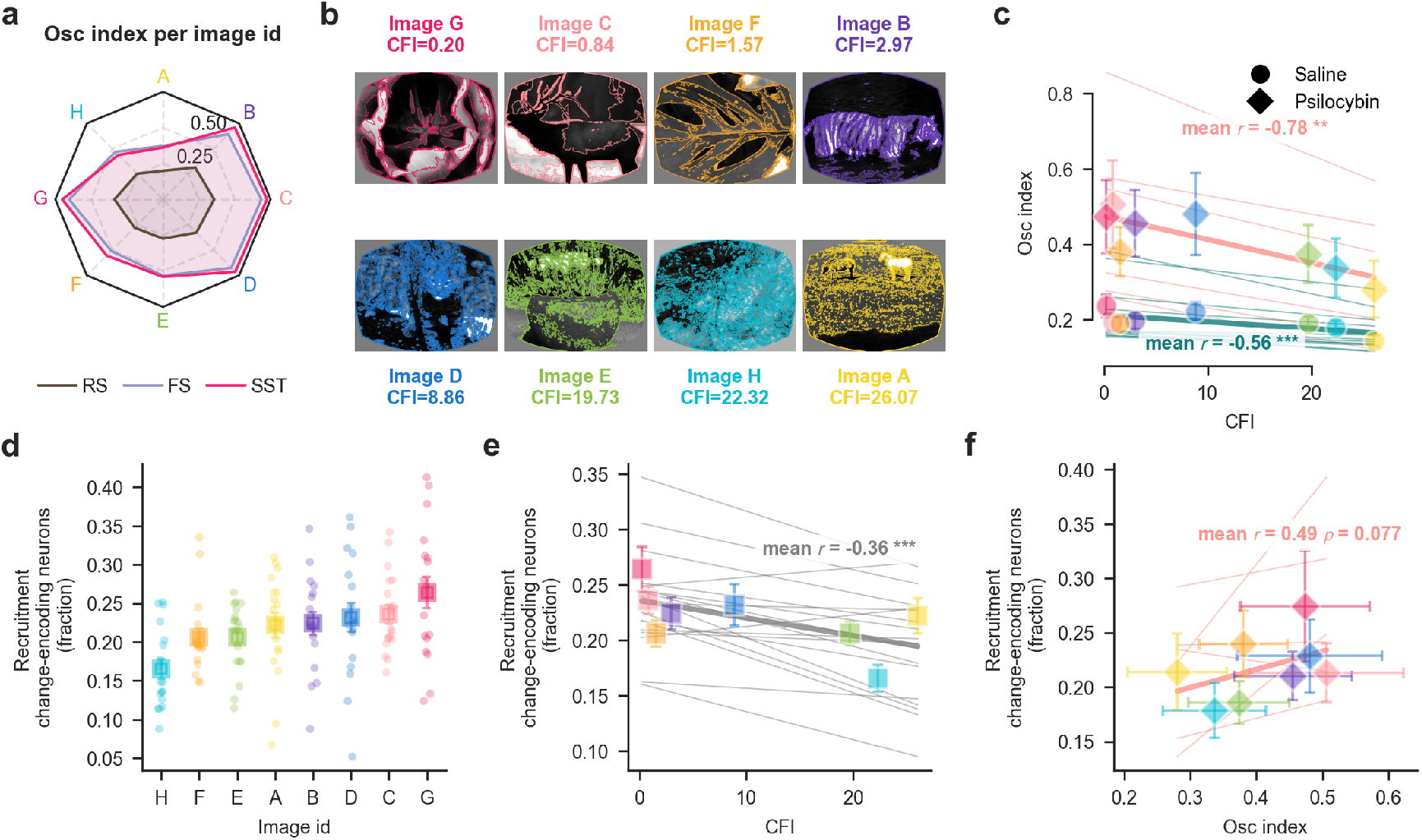
Stimulus features determine the magnitude of psilocybin-induced modulation. (a) Magnitude of psilocybin-induced modulation shown separately for each image A through H in the stimulus set. (b) The eight natural images used in the task. Colored overlays indicate detected contours used to compute the Contour Fragmentation Index (CFI). (c) Correlation between CFI and Osc index across images in psilocybin (diamonds) and saline (circles) sessions. Markers show means across sessions, error bars indicate s.e.m. and thin lines show individual-session fits. Reported r values are mean within-session Pearson correlations after Fisher transformation. P values were obtained using two-sided one-sample (t)-tests against zero. *P < 0.05, ***P < 0.01, ***P < 0.001. (d) Recruitment of change-encoding neurons following genuine image changes across all sessions, shown separately for each image. (e) Relationship between CFI and recruitment of change-encoding neurons across images across all sessions. (f) Relationship between Osc index and recruitment of change-encoding neurons across images in psilocybin (diamonds) and saline (circles) sessions. Error bars correspond to *±*1 s.e.m. Plotting and statistical conventions in E and F are as described for C.

We next asked whether this feature dependence reflected the tuning of the ensembles recruited by psilocybin. Change-encoding neurons were highly stimulus-selective, responding on average to only 1.86 *±* 0.04 (mean *±* s.e.m.) of the 8 images presented. Consistent with this sparse tuning, the fraction of change-encoding neurons recruited by genuine changes varied across stimuli (min-max range: 0.17 to 0.26; Figure 6d) and inversely correlated with the CFI (mean r=-0.36, P<0.001). Moreover, the magnitude of psilocybin-induced modulation showed a positive trend with the size of these image-specific change-encoding ensembles (mean r=0.49, P=0.077; Figure 6f). Thus, the image dependence of psilocybin-induced modulation may be related to the selective recruitment of feature-tuned change-encoding neurons.

## Discussion

Our behavioral results demonstrate that 1 mg/kg of psilocybin strongly impairs visual change detection in mice, with performance falling to chance. Using large-scale single-unit recordings, we found that this deficit was accompanied by a strong, visually-evoked 4-Hz oscillatory modulation, consistent with a recent report[24], and potentially reflecting altered thalamo-cortical interactions[25, 26]. This oscillation was strongest in change-encoding SST interneurons of the visual cortex, and transiently shifted population dynamics toward a state evoked by genuine image changes, that is, a surprising stimulus. Importantly, psilocybin did not trigger a uniform cortical state shift. Instead, this modulation remained constrained by the structural features of the visual input and was transient, mirroring the kinetics of the head-twitch response[27, 28], an established indicator of psychedelic potency in mice[29].

We propose that the shift toward surprise-like neural dynamics directly underlies the observed behavioral impairment. First, it is unlikely that the collapse in task performance stems from gross motor deficits, since in-depth video analyses did not reveal any clear impairment or alteration following psilocybin administration. Second, the simultaneous absence of both the behavioral and neural signatures in a single resilient mouse - neither showing a drop in performance nor a strong 4-Hz oscillation - within our cohort suggests that neural and behavioral effects are related. Third, the behavioral disruption in our visual task may be a consequence of the visual distortions observed in humans under psychedelics[30]; indeed, psilocybin is not known to selectively suppress task motivation or engagement[31, 32]. Finally, the behavioral disruption cannot be attributed to a loss of information about image identity. Basic visual representations were fully preserved: receptive fields remained unchanged, and image identity could be decoded from the population as accurately under psilocybin as under saline controls. Although a heightened internal bias toward change-signaling might seem paradoxical given the near-complete absence of task attempts, *i.e.*, reduced hits without an increase in false alarms, these observations can be reconciled. One possibility is that the aberrant recruitment of change-encoding populations during repeated images disrupts the stable context required for the learned change report, so that genuine changes no longer reliably trigger the learned lick response. We note, however, that a direct causal link between our observed neural signatures and behavioral performance remains to be established.

Current predictive processing models of acute psychedelic effects propose a disruption in the distinction between expected and unexpected sensory signals[4–6]. Our results provide a circuit mechanism for this phenomenon: psilocybin selectively amplifies change-encoding populations, causing even predicted, repeated images to evoke activity associated with contextual novelty. At the cellular level, explanations of psychedelic action have focused predominantly on deep–layer pyramidal neurons[4, 27, 33–35]. Our findings broaden this account by identifying a previously unrecognized effect of psilocybin on change-encoding SST interneurons: both the neural activity shift towards contextual novelty as well as the psilocybin–induced oscillation was strongest in SST interneurons, which have been implicated in encoding sensory-feedback discrepancies[9, 11, 36], navigation errors[10], and sensory adaptation[37–39]. Notably, serotonin, the natural ligand of the 5-HT2A receptor, has been associated with the broadcast of prediction errors and uncertainty[40–43]. Transcriptomic data reveal that some SST subtypes express 5–HT2A receptors[44–46], raising the possibility of direct modulation. While psilocybin inhibits SST neurons in the prefrontal cortex[47], psychedelic effects on these inhibitory networks may be highly region-specific or dependent on distinct cellular subtypes. Notably, the psilocybin-induced effects were strongest in visual cortical areas and absent in frontal cortical areas, with VISam showing the strongest relationship between change-encoding circuits and psilocybin-induced modulation. This suggests that higher visual areas are an important locus for psychedelic action.

This circuit-level account is consistent with reports that classical psychedelics reduce the neural distinction between predicted and unpredicted auditory and tactile stimuli in humans, as well as auditory stimuli in mice[48–50]. In rats, psychedelics made dopamine responses to predicted rewards resemble those to unexpected rewards during learning, consistent with increased prediction-error signaling[51]. Earlier behavioral studies in cats likewise reported that LSD reduced sensory adaptation[1, 52]. Thus, psychedelics may dismantle the adaptation strategy mice rely on to solve the visual change detection task[13]. More broadly, the disruption of operant performance in our mice parallels the suppression of classically conditioned freezing by psilocybin[53], consistent with psilocybin altering the neural representation of the conditioned stimulus and thereby weakening the associated fear response. Thus, psychedelics may act on a common predictive computation across sensory modalities and species, strengthening surprise signals.

Our findings provide insight into circuit mechanisms contributing to the therapeutic actions of psychedelics. Predictive-processing accounts propose that depression is characterized by overly precise priors that attenuate the influence of prediction errors and thus constrain belief updating[54–56]. The enhancement of surprise-related signals observed here could transiently oppose this imbalance. Given that SST dysfunction is a hallmark of depressive disorders[57–61], the prominent involvement of SST interneurons in the effects observed here may be particularly relevant to the therapeutic actions of psychedelics. A central open question is how the acute circuit reconfiguration observed here relates to the post-acute and longer-lasting effects of psychedelics. Recent work found reduced predictive suppression 24 h after psilocybin in mouse visual cortex as well as in recent human psychedelic users [62]. Psilocybin further induces lasting, 5-HT2A-dependent structural plasticity [63, 64], and brain-wide, region-specific modulation of neuronal bursting during the acute state [26] may provide an activity-dependent route for inducing long-term circuit changes. Beyond their potential clinical relevance, these acute alterations in contextual-novelty processing may provide a circuit-level explanation for the peculiar alterations in visual perception induced by psychedelics.

## Experimental procedures

### Data and code availability

The data from all experiments are available for download in Neurodata Without Borders (NWB) format on the DANDI Archive (https://dandiarchive.org/dandiset/001417). All custom code used to analyze this data is available at https://github.com/RobertoDF/psycode.

The following open-source software packages and libraries were used: NumPy, SciPy, scikit-image (skimage), Matplotlib, Pandas, xarray, scikit-learn, DeepLab-Cut, statsmodels, pingouin, iblatlas, keypoint-moseq, seaborn, joblib, NeMoS (https://github.com/flatironinstitute/nemos), spikeinterface[65], Jupyter (https://jupyter.org/), and pynwb (https://pynwb.readthedocs.io/en/stable/).

### Animals

All experiments were performed in Sst-IRES-Cre;Ai32(RCL-ChR2(H134R)_EYFP) mice (*n* = 10; 7 females and 3 males), aged 5–7 months. All procedures were approved by the Allen Institute’s Institutional Animal Care and Use Committee. After surgery, mice were single-housed and maintained on a reverse 12-h light cycle (20–22°C, 30–70% humidity), with experiments conducted during the dark cycle. For behavioral experiments, mice were water-restricted to maintain 85% of their initial body weight, with ad libitum access to food. The cohort consisted of 8 mice assigned to the drug treatment group (Day 1: sterile 0.9% saline; Day 2: psilocybin, 1 mg/kg) and 2 mice assigned to the control group (sterile 0.9% saline on both days). To avoid potential carryover from longer-lasting psilocybin effects, psilocybin was always administered on the second recording day. To control for possible effects of this fixed session order, two additional mice received saline on both days. All injections were administered intraperitoneally (i.p.), and Day 1 and Day 2 sessions were consecutive (24 h apart). This design yielded an initial potential pool of 12 saline sessions and 8 psilocybin sessions across the cohort. The psilocybin session for mouse 752309 was excluded entirely from both neural and behavioral analyses due to a synchronization error between the neural data, lick registrations, and visual stimulus timestamps. Task-performance analyses required a pre-injection perceptual sensitivity (AUC) *>* 0.6 on both recording days. Mouse 760322 did not meet this criterion (pre-injection AUC = 0.50 in both its saline and psilocybin sessions). This animal was excluded from the change-detection performance evaluation but was retained for all neural data analyses. Consequently, the behavioral analysis of change-detection performance (Figure 2b, c) included 11 saline sessions from 9 mice, and 6 psilocybin sessions from 6 mice. One saline session was excluded from the running-behavior analysis because of a sensor malfunction. For the analysis of neural activity, we excluded one outlier animal that showed no behavioral effect of psilocybin and continued performing the task maintaining a perceptual sensitivity above 0.6.

### Single-unit and local field potential recordings

Extracellular single-unit recordings and local field potential recordings using Neuropixels probes was conducted at the Allen Institute as part of the NIH-funded OpenScope project (https://alleninstitute.org/division/mindscope/openscope/). Mice underwent headplate implantation and craniotomy surgery. The cranial implant was a SHIELD implant described previously in detail[21]. Briefly, we created a CAD file of the implant with holes strategically placed above the areas of interest for our study. Hole positions were adjusted using Pinpoint[66]. Approximately a week after surgery, intrinsic signal imaging was used to identify visual area boundaries[67]. The mice were then habituated and trained in the behavior setups for several weeks and then to the Neuropixels rigs every day for a week. To prepare the brain for electrophysiological recordings, the coating over the implant was removed and replaced with a temporary layer of Kwik-Cast (World Precision Instruments). This was done a few days prior to recording to avoid using isoflurane the day of the experiment.

On the day of recording, the mouse was head-fixed, the protective well cap and Kwik-Cast layer were removed, and a layer of agarose or silicone oil was applied[68]. Six Neuropixels 1.0 probes[69] were targeted to visual areas (VISp, VISal, VISam, VISa), anterior cingulate (ACA), and primary somatosensory cortex (SSp–bfd). Probes were coated with CM–DiI (1 mM in ethanol; Thermo Fisher Scientific, V22888) for post–hoc localization. Each probe was mounted on a 3–axis micromanipulator (New Scale Technologies), aligned to its target opening, and manually lowered to the brain surface. After initial penetration (*∼* 100 *µ*m), probes were inserted automatically at 200 *µ*m min^-1^ to a final depth of *≤* 3.5 mm and allowed to settle for 15–30 min.

Data were acquired at 30 kHz (spike band, 500–Hz high–pass) and 2.5 kHz (LFP band, 1,000–Hz low–pass) using the Open Ephys GUI[70]. Videos of eye and body were acquired at 30 Hz; running wheel angular velocity was recorded at *∼* 60 Hz. Spike–band data were median–subtracted and processed with Kilosort4[71]. Quality labels were computed for each putative neuron using UnitRefine[72]. Putative double–counted spikes (spikeinter-face.curation.remove_duplicated_spikes) and artefactual units were removed (noise label by UnitRefine); remaining units were packaged into Neurodata Without Borders files.

We used published protocols to prepare whole mouse brains for clearing. Briefly, brains were per-fused and fixed in 4% paraformaldehyde for light-sheet microscopy. In a timeline of two weeks, the brain was stripped of lipids ([73]) and rendered transparent in an index matching solution ([73]), allowing for viewing the morphology of anatomical brain structures. Brains were then embedded in agarose for imaging ([73]). This protocol collection is ideal for experiments where it is necessary to preserve endogenous fluorescence. Agarose blocks containing cleared mouse brains were imaged by a light sheet microscope (LifeCanvas Technologies).

After the brains were processed in the imaging pipeline, Neuroglancer (https://Neuroglancer-demo.appspot.com) was used to reconstruct probe tracks in the brain. Points were placed along the length of the probe track and were closely inspected to ensure annotation of the probe tip and each track is assigned to a particular day of recording. Electrophysiological features recorded from neural probes were aligned with anatomical landmarks based on the Allen Mouse Brain Common Coordinate Framework (CCFv3). For this, we used the IBL alignment GUI (https://github.com/AllenNeuralDynamics/ibl-ephys-alignment-gui).

### Optotagging to identify SST interneurons

Following receptive field mapping and behavioral task (see below), the same optotagging procedure was performed on all mice. Blue light was delivered via a 465 nm LED (Plexon) controlled by a Cyclops LED driver or a 473 nm laser (Laser Quantum Ciel or Cobolt 06–MLD). The light source was coupled to a 400 µm diameter fiber optic cable (Thorlabs), with the tip positioned to illuminate the cranial window located in visual cortex. The following three stimulus types were presented at each of three light levels, with all nine combinations randomly interleaved: 6 ms pulses at 10 Hz, 10 ms pulses at 5 Hz, and a 1 s raised cosine ramp. Stimuli were presented at intervals of 1.5 s plus a uniformly distributed delay of 0–0.5 s. Mice had no prior exposure to the optotagging stimulus before the recording session. Optotagged cells were identified based on their responses to the 5 Hz stimulation. For each neuron, we examined each pulse individually. For every pulse, we calculated the firing rate in a 10-ms window beginning 2 ms after pulse onset and compared it to the number of spikes in a baseline window from –11 to –1 ms before pulse onset. A Wilcoxon signed–rank test was then applied across all pulses to determine whether post-stimulus and baseline spike counts differed significantly. Because some pulses yielded zero spike counts in both windows, we used a zero–handling method (zero_method=‘zsplit’ in scipy.stats.wilcoxon) that splits ties equally between positive and negative ranks, providing a balanced treatment of zero differences. Neurons with a significant difference (two-sided *P <* 0.05) were classified as SST neurons.

### Change detection task

Mice were trained in custom-designed, sound-attenuating behavior enclosures equipped with a 24 gamma-corrected LCD monitor (ASUS PA248Q). Head-fixed mice were positioned on a behavior stage with a 6.5 running wheel tilted upward by 10–15°. The monitor was placed 15 cm from the right eye, and visual stimuli were spherically warped to maintain constant perceived size, speed, and spatial frequency across the visual field. Water rewards were delivered via a solenoid (NI Research, 161K011) through a blunted 17-gauge hypodermic needle (Hamilton) positioned 2–3 mm from the animal’s mouth.

Mice were trained 1 h/day, 5 days/week on a go/no–go change detection task (Figure 1a). Mice learned to lick a reward spout when the identity of a briefly flashed visual stimulus changed. Correct responses within a post–change window (150–750 ms) triggered a water reward. Each session consisted of a continuous series of images. Change times were drawn from a truncated exponential distribution ranging from 2.25 s to 8.25 s (mean = 4.25 s) following sequence onset.

Each image presentation was treated as an individual trial. On each trial, the animal’s licking behavior was classified relative to the trial type (change vs. non–change). A hit was defined as a lick on a change trial in a window up to 0.9 s after image onset, a miss as no-lick on a change trial, a false alarm as a lick on a non–change trial in a window up to 0.9 s after image onset, and a correct rejection as no-lick on a non–change trial. False alarms were assigned only at least 4 images away from a change image to avoid miscategorization of reward-licks. For each session and each experimental epoch (e.g., pre–injection, post–injection), the proportion of each outcome was calculated separately for change image and no–change image trials. Specifically, hit and miss proportions were computed within change trials (hit rate + miss rate = 1), and false alarm and correct rejection proportions within non–change trials (false alarm rate + correct rejection rate = 1). To assess the effect of the drug manipulation on behavioral performance, a two–way repeated–measures analysis of variance (RM–ANOVA) was conducted separately for hit rates (image change trials) and false alarm rates (no–change trials). The within–subject factors were condition (saline vs. psilocybin) and epoch (pre–injection vs. post–injection). The dependent variable was the proportion of hits or false alarms per session. The analysis was performed using the rm_anova function from the Pingouin library. The main statistic of interest was the condition *×* epoch interaction effect, which indicates whether the change in performance from pre– to post–injection differed between drug conditions.

### Motion sequencing analysis

Body part tracking was performed with DeepLabCut (version 3.0.0rc10)[74]. We labeled 109 frames from a frontal video and 99 frames from a lateral video for training (both 658 *×* 792 pixels, 60 fps). The following keypoints (body features) at the frontal camera were defined: frontpaw left, frontpaw right, hindpaw left, nose base, nose tip, ear base, ear mid. For the lateral camera, we used the following keypoints: frontpaw left, frontpaw right, hindpaw left, hindpaw right, tail base, tail mid, tail tip, nose tip, ear tip. 95% of frames were used for training a ResNet–101 network (default parameters, 200 iterations, 5 shuffles) with the following parameters: Training error, 1.06 pixels (frontal) and 2.23 pixels (lateral); test error, 5.7 pixels (frontal) and 4.6 pixels (lateral); cutoff *P* -value = 0.1 for both *x* (medio-lateral for frontal camera, antero-posterior for lateral camera) and *y* (ventro-dorsal) coordinates before modeling.

Unsupervised behavioral classification used Keypoint–MoSeq (v0.6.7)[16]. Outlier keypoints were removed (scale factor = 6.0). PCA on aligned, centered keypoints selected 5 latent dimensions (minimum explaining 90% variance). *κ* was tuned to 10^6^ (autoregressive–only model) and 10^5^ (full model) to achieve a median syllable duration of more than 10 frames. After fitting the AR–HMM, the model was reapplied to training and new data. Syllable frequency and duration were analyzed with MoSeq’s function ‘plot_syll_stats_with_sem‘ setting a minimum syllable frequency of 0.01 and restricting the analysis to post-injection epochs.

### Single-unit analysis

Following automated spike sorting, units were classified using UnitRefine[72], an automated curation framework trained to reproduce expert labels by integrating complementary features of spike timing, waveform morphology, amplitude stability, and cluster isolation. UnitRefine assigned each cluster to single-unit activity (SUA), multi-unit activity (MUA), or noise. Units were retained if they were classified as SUA or, if they met the following quality metrics: ISI violations *<* 0.5 (the relative firing rate of contaminating spikes, estimated from interspike intervals shorter than 1.5 ms and normalized by the overall spike rate), amplitude cutoff *<* 0.1 (an estimate of the fraction of spikes missed because their amplitudes fell below the detection threshold, calculated from the symmetry of the spike-amplitude distribution), and presence ratio *>* 0.9 (the fraction of 100 equal-duration recording blocks containing at least one spike from the unit; values above 0.9 indicate detection in more than 90% of the session). Only units with a firing rate of at least 1 spike/s in both the pre-and post-injection behavioral epochs were included. These thresholds ensured well–isolated, stable units with minimal contamination from multi–unit activity. Optotagged SST interneurons were included regardless of these thresholds, as their identity was independently confirmed by light–evoked responses. Fast–spiking (FS) neurons were identified based on a trough–to–peak latency *<* 0.4 ms.

For population–level comparisons across conditions, additional inclusion criteria were applied to ensure robust statistical power. For RS and FS neurons, cortical areas were included only if they contributed at least three sessions per condition (saline and psilocybin), with each session containing at least seven neurons of that type. For the sparser SST population, areas were included if they had at least three sessions per condition with a minimum of four neurons per session. These thresholds balanced statistical power with the retention of as many areas as possible given the lower yield of interneurons.

For each isolated unit, spike density functions (SDFs) were computed across a temporal window of -1.0 to 1.5 s relative to either the image change or all repeated, no-change image presentations. Spike trains were binned at 1 ms resolution and convolved along the temporal axis with a causal, right-aligned exponential filter window:

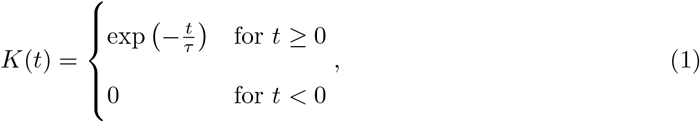

where the decay time constant was set to *τ* = 5 ms. The resulting convolved arrays were divided by the sampling interval to convert counts to firing rates (spikes/s). To normalize firing rates, z–scoring was performed using baseline statistics (mean and standard deviation) computed from the entire pre–injection change–detection block, the full behavioral session before drug administration.

This extended window provided stable estimates even for neurons with extremely sparse firing. The same pre–injection statistics were used to normalize spike counts during all stimulus periods (both pre– and post–injection), expressing each neuron’s responses relative to its own pre–injection baseline.

### Behavior-adjusted firing rates

To quantify neural activity not explained by ongoing motor behavior, we fitted a generalized linear model (GLM) for each neuron using the ‘PopulationGLM‘ implementation from the NeMoS library (https://github.com/flatironinstitute/nemos). Behavioral predictors included the first 60 FaceMap motion components from each of three video-derived streams (face, whisker region, and body) yielding 180 video-derived predictors per session[20]. The whisker region was analyzed separately because motion energy in the full-face video was dominated by paw movements, allowing whisker-specific motion to be captured more precisely. Running speed and pupil area were also included as predictors, each represented using five raised-cosine basis functions. Predictors were averaged within 1-s bins, aligned to the neural data, and standardized. Stimulus regressors were intentionally omitted because visual stimuli were identical across pre-and post-injection epochs and across psilocybin and saline groups. Thus, any stimulus-driven neural activity was present in both conditions and was not expected to confound comparisons between treatment groups. The aim of the model was not to maximize predictive accuracy, but to isolate neural activity that could not be explained by the measured behavioral variables.

Spike counts were calculated in 1-s bins, and only neurons with a mean pre-injection firing rate greater than 1 Hz were included. The model was fitted using data ending 120 s before injection and was then used to predict firing rates from behavioral measurements beginning 120 s after injection. The peri-injection interval was excluded to minimize contamination from neural and behavioral responses associated with experimenter handling. Each neuron had its own set of model coefficients, although neurons were processed in batches of 100 for computational efficiency. Models used a Poisson observation model with an exponential inverse-link function and Lasso regularization (regularization strength = 0.01). For neuron *i* and time bin *t*, the Pearson residual was calculated as

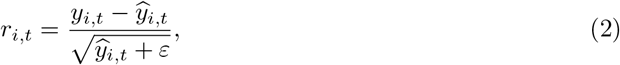

where *y_i,t_* is the observed spike count and *y_i,t_* is the spike count predicted by the GLM. The constant *ε* prevented division by zero when predicted values approached zero. Negative residuals indicated less firing than predicted from behavior, whereas positive residuals indicated greater firing than predicted.

Pearson residuals for individual neurons were smoothed using a centered 120-s rolling mean and sampled at 30-s intervals. For each session, brain area, and neuron type (RS, FS, SST), we then calculated the median residual across the available neurons, providing an outlier-robust session-level estimate of neural activity unexplained by the modeled behavioral variables.

### Oscillation index

To quantify the psilocybin-induced oscillatory modulation on a single-neuron level, we calculated the ΔPSTH as the difference between the post-injection and pre-injection peri-stimulus time histograms, z-scored for each individual neuron. After smoothing each ΔPSTH with a centered 50-ms Gaussian window (SD = 10 ms), a cosine function was fitted to its ΔPSTH over the interval of 0–750 ms post-stimulus onset using nonlinear least-squares optimization (Levenberg-Marquardt algorithm, implemented via scipy.optimize.curve_fit). The model was defined as:

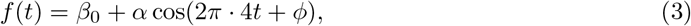

where *β*_0_ is the baseline offset, *α* is the modulation amplitude, and *ϕ* is the phase. To ensure a standardized representation where the amplitude is strictly positive, if a negative *α* was returned, we inverted its sign and advanced the phase by *π* (*ϕ → ϕ* + *π*). We then evaluated the model residuals as the difference between the empirical ΔPSTH and the fitted curve *f* (*t*). Fit quality *Q* was defined as the ratio of the amplitude to the standard deviation of these residuals:

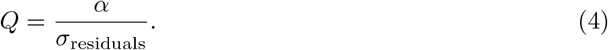

We then defined the Osc index as the fitted amplitude multiplied by a logistic weight determined by *Q*:

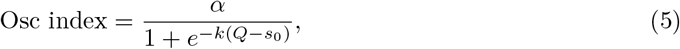

with *s*_0_ = 0.5 and *k* = 15. These parameters produce a steep transition around *Q* = 0.5, strongly attenuating the amplitudes of fits whose residual variability is large relative to the fitted oscillatory component while preserving the amplitudes of well-fitting 4-Hz oscillations. Thus, the Osc index represents the magnitude of the fitted 4-Hz modulation weighted by its signal-to-residual ratio. When comparing saline and psilocybin groups we further verified that the rhythmic modulation reflected a sustained oscillation rather than only modulation of the image onset and offset responses, we separately analyzed the late-phase window from 500 to 750 ms, corresponding to the third oscillatory peak. This segment was repeated three times to generate a 750-ms trace, to which the same model was fitted to obtain the late-Osc index. Neurons were classified as oscillatory (‘Osc neurons’) if both their Osc index and late-Osc index exceeded a threshold defined as the 80th percentile of the Osc index across all cortical neurons in the psilocybin condition.

### Mutual information analysis

To quantify the information each neuron carried about image identity and image change, we computed mutual information (MI) between spike counts of the respective categorical variables. For image change encoding, the target variable was a binary vector indicating whether a trial was a change trial (image identity different from the previous reference) or a non–change trial (repetition of the reference image). For image identity encoding, the target variable was the specific image identity (image id 1–8). Both MI values were computed using the mutual_info_classif function from scikit–learn[75], which is appropriate for discrete features and targets.

Spike counts were extracted per image presentation from both pre and post–injection epochs. For change encoding, we used the early post–stimulus window (0–100 ms after image onset) to capture the rapid detection of a deviation while minimizing potential contamination from reward–related activity. For identity encoding, we used the full stimulus window (0–250 ms after image onset) to capture the sustained representation of the natural image.

Neurons were classified as image change–encoding or image id–encoding if their MI value fell above the 80th percentile of the respective distribution across the population (cf. Figure S5). This threshold was calculated focusing on areas with significant visual information encoding (VISa, VISal, VISam, VISp).

### Receptive field analysis

Each experiment included two receptive-field mapping blocks, one before and one after injection. Drifting Gabor patches (2 Hz, 0.04 cycles/degree) were presented within a 20° circular mask at 81 randomly ordered screen locations arranged in a 9 *×* 9 grid. Each stimulus was presented for 250 ms without an intervening blank period.

For each unit, spatial receptive fields (RFs) were calculated separately for the pre-and post-injection mapping blocks. Spike counts were measured during a response window extending from stimulus onset to 0.24 s after onset. This window was 10 ms shorter than the 0.25-s stimulus duration to avoid including spikes associated with the immediately subsequent stimulus. For each spatial coordinate (*x, y*), the mean firing rate was calculated as

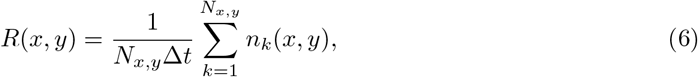

where *N_x,y_* is the number of presentations at coordinate (*x, y*), *n_k_*(*x, y*) is the spike count during presentation *k*, and Δ*t* = 0.24 s. The resulting 9 *×* 9 matrix contained the mean response rate at each stimulus position.

Before permutation testing, units were screened using an extremity index calculated from the complete spatial response matrix:

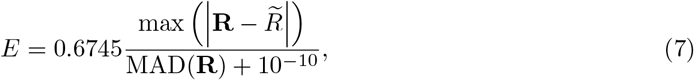

where **R** is the vector of responses across all spatial positions, *R* is its median, and MAD(**R**) = median(*|***R** *− R|*). When the median absolute deviation was zero, the extremity index was set to zero. Only units with *E >* 4 were submitted to permutation testing.

Statistical significance was evaluated using circular shuffling of spike times. Spike times were first restricted to the duration of the corresponding receptive-field mapping block. For each shuffle, the complete spike train was shifted by a uniformly sampled temporal offset and wrapped around the boundaries of the mapping block. This procedure preserved the number of spikes and their relative temporal structure while disrupting their alignment to stimulus presentations. A shuffled spatial response matrix was then calculated using the same procedure as for the observed data.

For each spatial position, a two-sided permutation *P* -value was calculated as

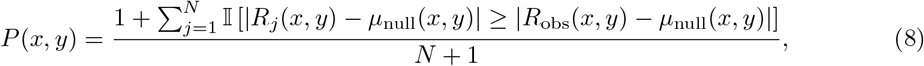

where *R*_obs_(*x, y*) is the observed response, *R_j_*(*x, y*) is the response obtained in shuffle *j*, *µ*_null_(*x, y*) is the mean shuffled response, and *N* is the number of shuffles. The 81 spatial *P* -values were corrected using the Benjamini–Hochberg false-discovery-rate procedure.

Permutation testing was performed in two stages. An initial screening stage used 800 shuffles and an FDR-corrected threshold of *α* = 0.15. Units without any significant spatial positions at this stage were excluded from subsequent fitting. Retained units were then evaluated using 8,000 shuffles and an FDR-corrected threshold of *α* = 0.05, producing the final binary significance mask. The final significance mask was convolved with an eight-connected 3 *×* 3 neighborhood kernel:

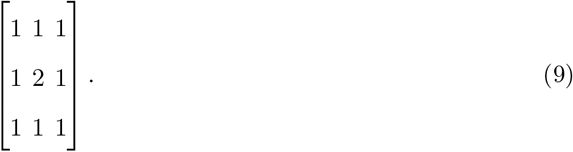

The empirical RF center was defined as the spatial coordinate at which this convolved mask reached its maximum. When multiple positions shared the maximum, their row and column indices were averaged and rounded to the nearest grid position. RF width was quantified by convolving the binary significance mask with an eight-connected neighborhood kernel and taking the maximum value of the resulting matrix. This provided a discrete measure of the local spatial extent of significant RF responses.

Response sign was determined from the mean z-scored response within the final significance mask. Units with a mean above zero were classified as having positive RFs, whereas those with a mean below zero were classified as having negative RFs.

For retained units, a rotated two-dimensional Gaussian model was fitted to the complete raw spatial response matrix:

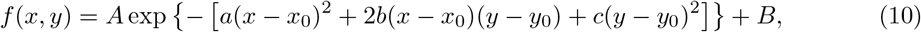

with

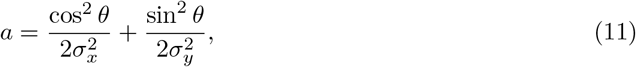

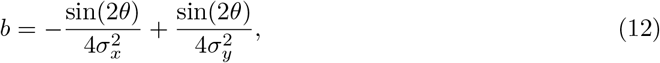

and

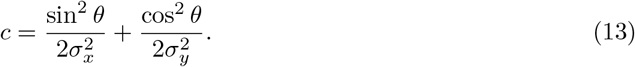

Here, *A* is the response amplitude, *B* is the offset, (*x*_0_, *y*_0_) is the fitted center, (*σ_x_, σ_y_*) are the spatial standard deviations along the two Gaussian axes, and *θ* is the rotation angle. The empirical RF center obtained from the convolved significance mask was used to initialize (*x*_0_, *y*_0_). Initial width estimates were calculated from absolute-response-weighted second moments around this center and were constrained to be at least one-half of the spatial grid spacing. The initial rotation was 0*^◦^*. For positive RFs, *A* was constrained to be non-negative; for negative RFs, it was constrained to be non-positive. Center coordinates were bounded by the sampled spatial range, *σ_x_* and *σ_y_* were bounded between one-half of the corresponding grid spacing and 50*^◦^*, and *θ* was bounded between *−*90*^◦^*and 90*^◦^*.

Parameters were estimated using bounded nonlinear least squares with the Trust Region Reflective algorithm and a soft-L1 loss function. If optimization failed, the initial parameter values were retained. Fits with an RF-width value *≤* 2 were subsequently invalidated by setting their Gaussian parameters and goodness-of-fit values to missing values.

Goodness of fit was quantified using the adjusted coefficient of determination:

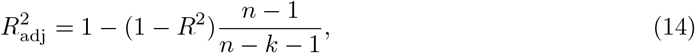

where *n* = 81 is the number of spatial positions and *k* = 7 is the number of fitted parameters. Additional measurements included the fitted center (*x*_0_, *y*_0_), *σ_x_*, *σ_y_*, orientation, aspect ratio,

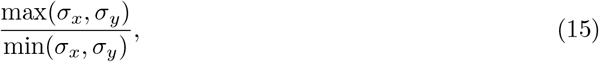

and ellipticity,

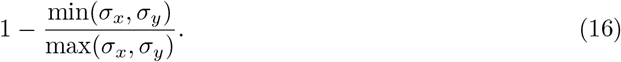

The extremity index and the mean, minimum, and maximum response rates across the complete spatial response matrix were also retained for each unit.

### Population decoding

To investigate how neural ensembles represent task variables and visual features across drug conditions, we used a population decoding pipeline. We constructed balanced-class linear classifiers applied to spike counts starting from stimulus onset over progressively longer windows. Spike counts were integrated across a grid of temporal parameters with window durations ranging from 10 to 300 ms in 10-ms increments, beginning at stimulus onset. Population decoding was performed using a logistic regression model implemented via scikit-learn. To account for unequal trial distribution across decoded classes, class weights were inversely proportional to class frequencies:

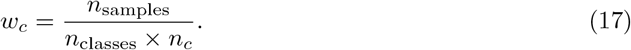

Feature matrices were normalized (z-scored) across trials before model fitting. The pipeline decoded two primary target variables. For image change decoding, a binary classification determined whether a stimulus presentation constituted a change event. For stimulus identity, a multiclass classification identified the specific visual image presented. Models were evaluated using two distinct cross-validation strategies to assess the stability of neural representations. The within-block decoding strategy handled single-epoch decoding by utilizing a stratified k-fold cross-validation scheme where *k* = 5 to approximately preserve class proportions in each fold. Alternatively, the cross-block decoding strategy assessed the stability of representations across behavior by training models on data from the baseline pre-injection epoch and testing on data from the post-injection epoch. To directly quantify the stability of the population code across these epochs, we calculated the ‘generalization Δ’, defined as the drop in decoding accuracy between the baseline within-block decoder and the cross-block decoder. Performance for both strategies was quantified using balanced decoding accuracy to guard against any residual class imbalances. To evaluate statistical differences in the temporal trajectory of decoding performance between saline and drug conditions, independent two-sample t-tests (assuming equal variance) were performed to compare generalization Δ values between the saline and psilocybin injection groups within each temporal window. To control for the inflation of type I errors resulting from family-wise testing across the time-window grid, the set of p-values across all unique temporal windows was adjusted for multiple comparisons using the Benjamini-Hochberg false discovery rate (FDR) procedure.

### Population trajectory analysis

To assess whether psilocybin reconfigured population-level image or image change representations, we analyzed the time-varying activity of simultaneously recorded neural populations across experimental epochs (pre- versus post-injection), drug conditions (saline versus psilocybin), and trial types (change versus no-change). For each session, cell type, and condition, PSTHs were constructed by aligning spiking activity to image onset over a window of 0 to 100 ms, and z-scored on a per-neuron basis. Only sessions with at least 20 neurons of a given cell type were retained for this analysis. For visualization purposes across this 0 to 100 ms epoch, we built a population matrix by concatenating all neurons from all sessions of a given condition and cell type. To capture the dominant response patterns in our neural ensembles, we applied PCA based on the pre-injection data (change and no-change trials combined). The first three PCs were used to project all conditions into a shared low-dimensional space (Figure S11). This projection provides an intuitive view of the population trajectories. The quantitative analysis was performed in the full, neuron-dimensional space without dimensionality reduction, using the z-scored PSTHs directly. For each session and cell type, we defined two time-resolved difference vectors:

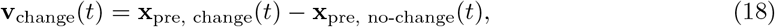

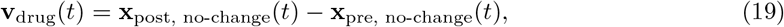

where **x**(*t*) represents the population vector (with a length equal to the number of neurons) at time *t*. At each time point, we computed the scalar projection of the drug-effect vector onto the direction of the image-change vector:

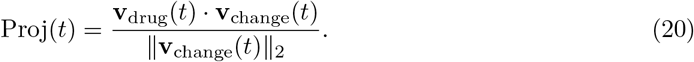

This projection measures how strongly, and in which direction, the drug shifts the population along the endogenous image-change axis. To obtain a single score per session, the time-resolved projection was averaged over an analysis window of 20 to 80 ms after image onset. Finally, this integrated score was divided by the number of neurons in that session to yield the mean projection per neuron (expressed in arbitrary units, denoted as “Change-alignment (a.u.)”), making sessions with different neuron yields directly comparable. For psilocybin sessions, a null distribution was generated via a temporal shuffle procedure. For each shuffle iteration (1,000 iterations), the time indices of the complete drug-effect trajectory **v**_drug_(*t*) were randomly permuted across the entire PSTH epoch prior to window selection, leaving the visual-change trajectory **v**_change_(*t*) intact. This procedure breaks the true time-resolved relationship between the two trajectories while fully preserving the underlying population structure, cell identities, and response amplitudes. The mean projection per neuron was then recomputed for each iteration over the same 20 to 80 ms analysis window. The average across these shuffle iterations defined the session-wise chance level. Statistical significance of the difference between the saline and psilocybin cohorts was assessed using a two-sample permutation test (20,000 iterations) on the per-session mean projection per neuron, performed independently for each cell type.

### Active fraction analysis

To examine whether psilocybin modulated the recruitment of functional ensembles (either change-or identity-encoding) across sequential trial presentations, we calculated the active fraction of image id-encoding and image change-encoding neurons, defined as the proportion of neurons within an ensemble that were significantly activated on a single-trial basis. For image change-encoding ensembles, candidate neurons were initially selected if the MI between their spike counts and the trial type (change versus no-change) exceeded the 70th percentile of the population distribution. To ensure strict image specificity, these candidate units were then subjected to a secondary, more stringent binomial test (*P <* 0.05) paired with a minimum absolute response difference constraint of 10 spikes/s within a 0-100 ms post-stimulus window. Specifically, we tested whether a neuron’s response on a change trial with a given image was significantly greater than its response on no-change trials where that same image was repeated. Neurons meeting these dual criteria were classified as dedicated change-encoding cells for that specific image identity, capturing the localized tuning of change responses. Conversely, for image id-encoding ensembles, this secondary image-specific filtering was omitted. Instead, selection was restricted to neurons that uniquely encoded image identity, explicitly excluding any units exhibiting dual encoding to both identity and change. Neuronal responsiveness was determined on a trial-by-trial basis for each image presentation by quantifying spike counts within the 0-100 ms window following image onset. For each neuron-image pair, an initial baseline activation threshold (*θ*_pre_) was established using pre-injection no-change trials:

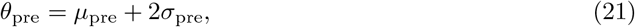

where *µ*_pre_ and *σ*_pre_ represent the mean and standard deviation of the post-stimulus spike count across pre-injection no-change trials, respectively. To account for state-dependent fluctuations or baseline firing rate drift over the course of an extended recording session, as well as potential psilocybin-induced changes in baseline firing rates, post-injection thresholds were adjusted dynamically: the baseline firing rate for each neuron (*µ*_baseline_) was calculated within a 250 ms pre-stimulus window, evaluated separately for the pre-and post-injection blocks. The difference between these pre-stimulus baselines was then added to the initial threshold to yield a drift-corrected post-injection threshold (*θ*_post_):

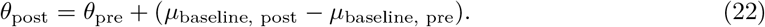

A neuron was classified as active on any given trial if its post-stimulus spike count exceeded its corresponding image-specific, drift-corrected threshold (*θ*_pre_ for pre-injection trials and *θ*_post_ for post-injection trials). The active fraction was computed trial-by-trial as the proportion of responsive neurons within the target ensemble. This metric was calculated separately for the pre-and post-injection epochs to generate a continuous time series aligned to the moment of injection. Population-level time courses were constructed by aggregating these time series across sessions. For each temporal bin, the mean active fraction was compared between the saline and psilocybin conditions using an independent-samples, two-tailed t-test. Normality was verified for the majority of bins; in instances where the normality assumption was violated, statistical significance was confirmed using non-parametric Mann-Whitney-U tests.

### Image contour analysis

To quantify the contours of the natural images used in the change-detection task, we developed a Contour Fragmentation Index (CFI). This metric captures the degree to which an image is composed of numerous, short, or disjointed edges, which we hypothesized would scale with the recruitment of change-signaling ensembles. The CFI is calculated as the ratio of the total number of discrete contours (*N*) to their average spatial length (*L̄*):

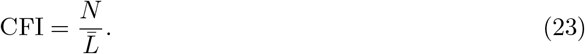

We implemented a multi-threshold framework to extract contours across diverse luminance levels. Images were first smoothed using a Gaussian filter (*σ* = 2.0 pixels) to mitigate high-frequency noise. We then determined a set of optimal intensity thresholds using the Multi-Otsu algorithm configured for four classes. For each identified threshold, contours were extracted from both the original luminance maps and their inverted negative images via the marching squares algorithm (skimage.measure.find_contours). This dual-polarity approach ensured that the index comprehensively accounted for both light-on-dark and dark-on-light structural elements across the full dynamic range of the stimulus. The resulting CFI provides a scalar measurement of image fragmentation; a higher CFI indicates a complex composition dominated by many small, disjointed features, whereas a lower CFI tracks an image with fewer, more continuous structural elements. We then used linear regression to evaluate the relationship between the CFI of each stimulus and its corresponding magnitude of psilocybin-induced neural modulation.

To assess relationships across images, neural measures were first averaged across neurons within each session and image. Pearson correlations were then calculated across the eight images separately for each session. Correlation coefficients were Fisher-*z*-transformed and tested against zero using a two-sided one-sample *t*-test across sessions. The reported mean correlation was obtained by back-transforming the mean Fisher-*z* value. Mean within-session variance accounted for was calculated by squaring each session’s correlation coefficient and averaging the resulting *R*^2^ values across sessions.

### Statistical analysis

Statistical analyses were performed in Python (version 3.11) using the scipy.stats, statsmodels, and pingouin libraries. All statistical tests were two-sided unless noted otherwise, and the baseline significance threshold for all analyses was set at *α* = 0.05. Statistical significance is denoted as follows: *(*P <* 0.05), **(*P <* 0.01), and ***(*P <* 0.001).

Hierarchical bootstrap: To account for the nested structure of the neural recordings, where hundreds of individual neurons were sampled across a smaller cohort of sessions, we implemented a hierarchical bootstrapping approach. This framework avoids the inflation of statistical significance that occurs when treating individual cells as independent samples. For each comparison, we executed 10,000 bootstrap iterations. On each iteration, experimental sessions were first resampled with replacement, and the constituent neurons within those selected sessions were subsequently resampled with replacement. This bootstrap distribution was used to derive the population mean and the standard error of the mean (s.e.m.). Statistical significance was assessed using a two-sided bootstrap test. For each bootstrap iteration *i*, Δ*_i_* was defined as the difference between the bootstrapped condition means:

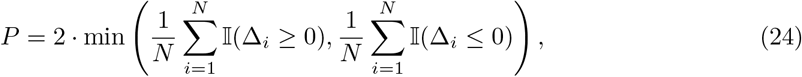

Here, *N* = 10,000 is the total number of bootstrap iterations, Δ*_i_* is the difference between the bootstrapped means of the two conditions in iteration *i*, and I(*·*) is an indicator function that equals 1 when its argument is true and 0 otherwise. *P* -values were bounded below by 1*/N*, yielding a minimum possible value of 0.0001.

Standard statistical tests and normality: For datasets where hierarchical nesting was not applicable, such as mouse-level behavioral parameters in Figure 1 and Figure 6, an automated analysis pipeline determined the most appropriate frequentist test. The normality of each group was evaluated using the Shapiro-Wilk test. For normally distributed data, parametric independent or paired t-tests were implemented, applying Welch’s correction in cases of unequal variance between independent cohorts. When the assumption of normality was violated, non-parametric alternatives were used, specifically the Mann-Whitney-U test for independent samples and the Wilcoxon signed-rank test for paired designs.

## Supplementary Information

**Fig. S1.**
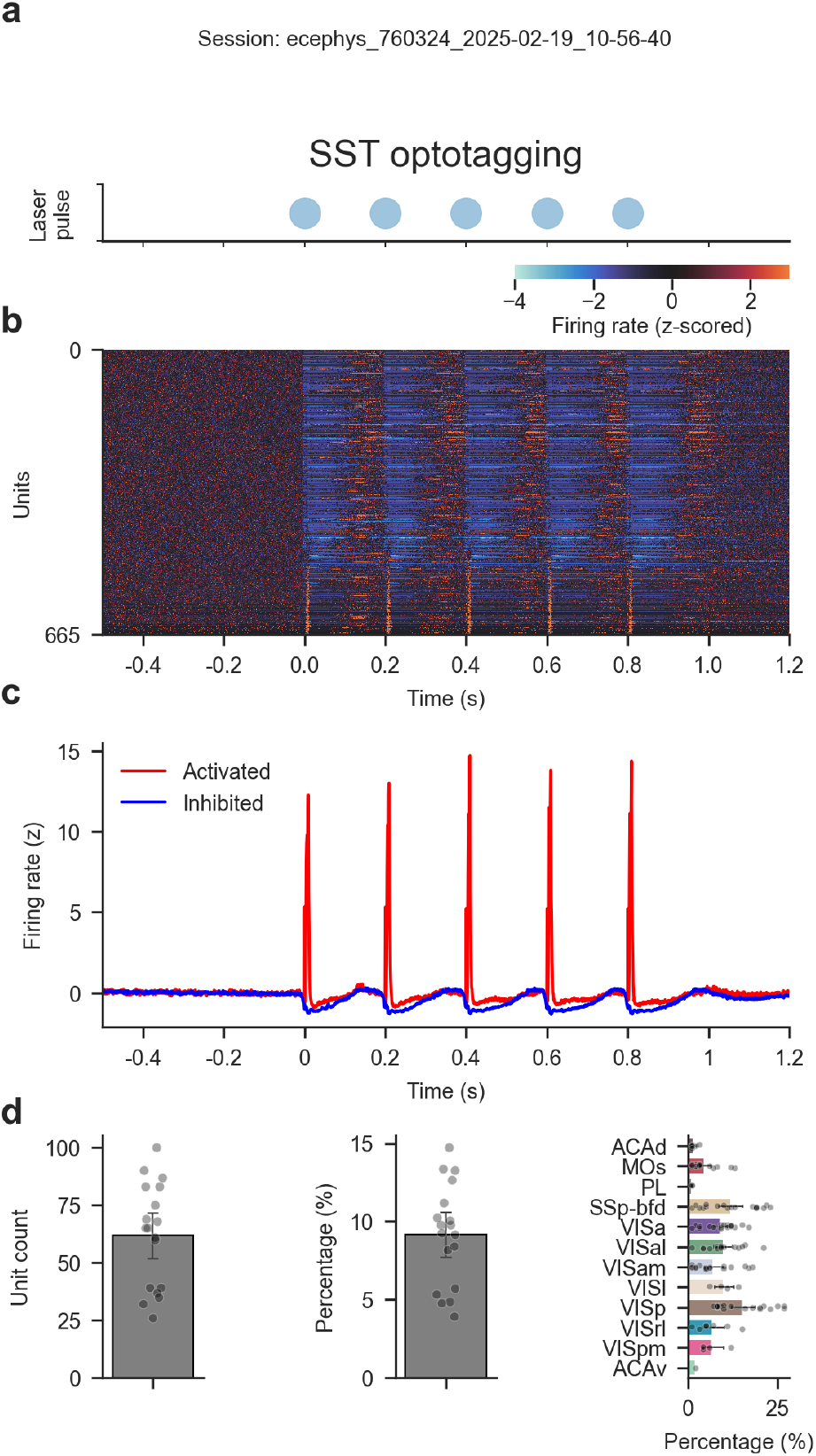
Optotagging identifies SST interneurons. (a) 5 Hz laser pulse used for the optotagging protocol aligned to (b) and (c). (b) Single-unit firing rates of an example session (ecephys-760324-2025-02-19-10-56-40). Shown here are all neurons with significant modulation following laser stimulation. (c) Average firing rate of activated and inhibited neurons in the same session as (b). (d) Average count (left) and percentage (center) of optotagged cortical neurons per session. Right: average percentage of optotagged neurons per cortical area.

**Fig. S2.**
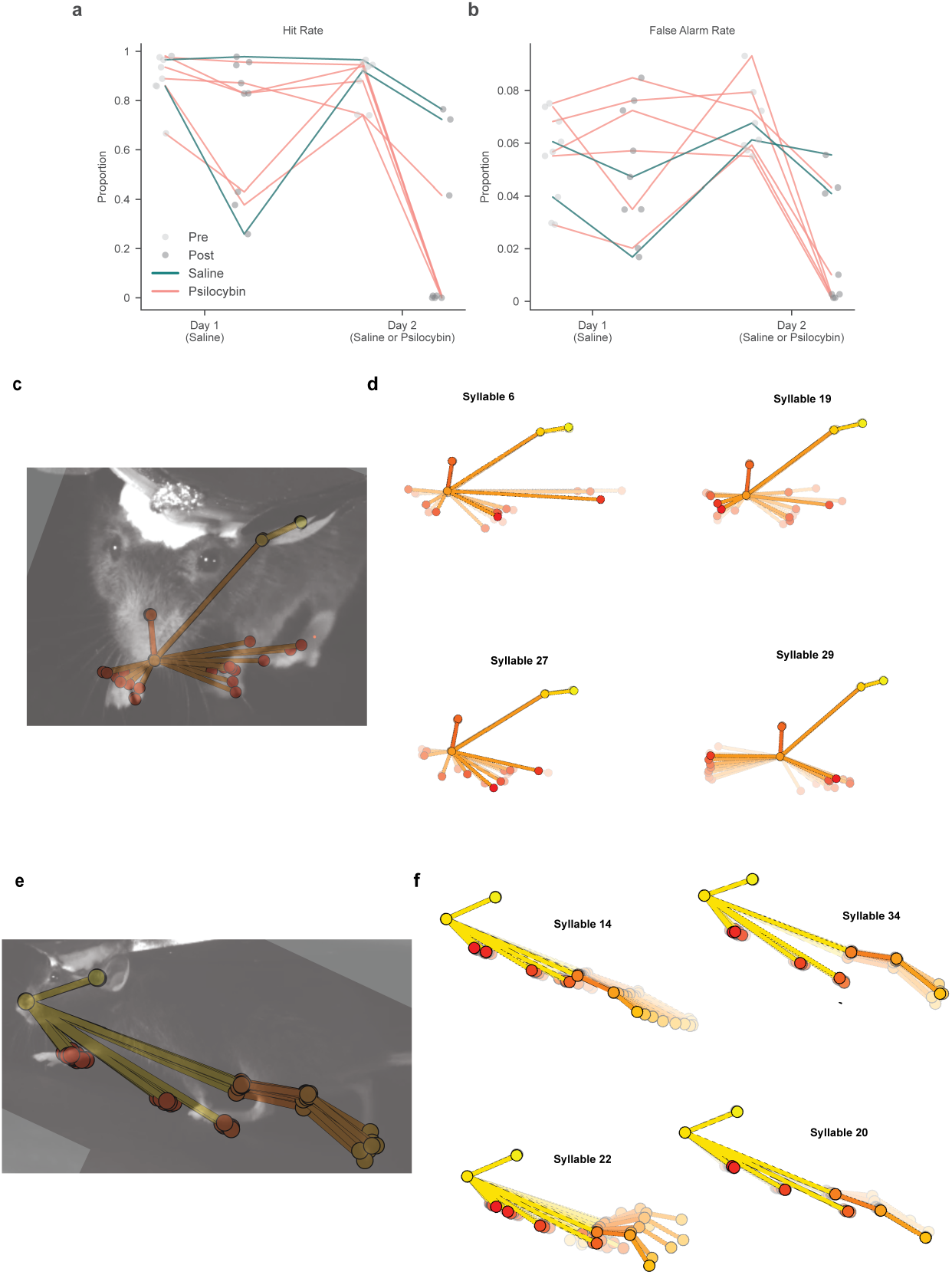
Behavioral performance and MoSeq analysis. (a) Hit rate measured before (Pre, light gray circles) and after (Post, dark gray circles) injection across Day 1 and Day 2. Each circle represents a single behavioral session. Lines connect the performance trajectories of individual animals across experimental epochs and days. Teal lines track control animals that received saline injections on both days (n = 2). (b) False alarm rate across experimental epochs and days, formatted identically to (a). (c) Frontal camera view with superimposed keypoints extracted with DeepLabCut. (d) Example syllables extracted from frontal camera videos. Each trajectory plot shows a sequence of poses along the average trajectory through latent space associated with a given syllable. (e) Same as (c) for lateral camera videos. (f) Same as (d) for lateral camera videos.

**Fig. S3.**
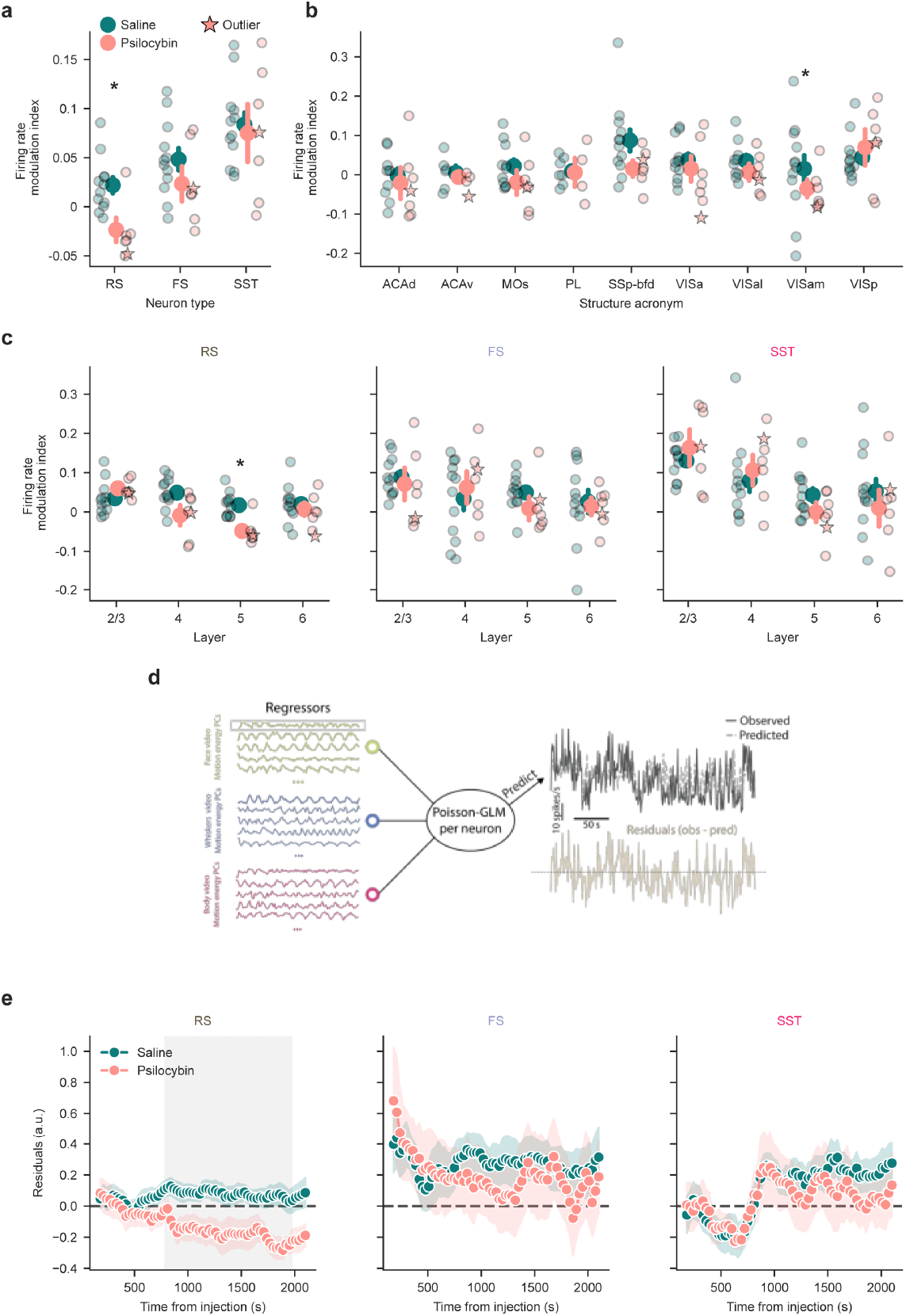
Psilocybin decreases activity in cortical RS neurons. (a) Firing rate modulation index across neuron types. Psilocybin causes a significant decrease of firing rates in RS neurons (psilocybin = -0.024 *±* 0.012, saline = 0.022 *±* 0.008, mean *±* s.e.m. across sessions, p = 0.048, hierarchical bootstrap). (b) Firing rate modulation index across brain areas, with no significant differences. (c) Firing rate modulation index across layers and neuron types. Psilocybin causes a significant decrease of firing rates in layer 5 RS neurons (psilocybin = -0.048 *±* 0.014, saline = 0.017 *±* 0.008, mean *±* s.e.m. across sessions, p = 0.019, hierarchical bootstrap). (d) Schematic of the GLM-based prediction. We extracted 40 principal components (PCs) from three behavioral videos using FaceMap. These PCs were used as regressors in a Poisson GLM fitted to each neuron’s activity during the pre-injection block. The model was then used to predict firing rates in the post-injection block. Residuals were computed as (observed - predicted) /√(predicted), yielding standardized residuals. (e) Grand average time course of residuals per group (psilocybin vs. saline). Residuals were first aggregated within each session using a median estimator (see Methods). RS neurons exhibit a significant decrease in firing rate beginning ∼30 min post-injection. Gray rectangle represents p <0.05 using Mann-Whitney-U test. Shading indicates *±*1 s.e.m. Error bars in all panels represent *±*1 s.e.m.

**Fig. S4.**
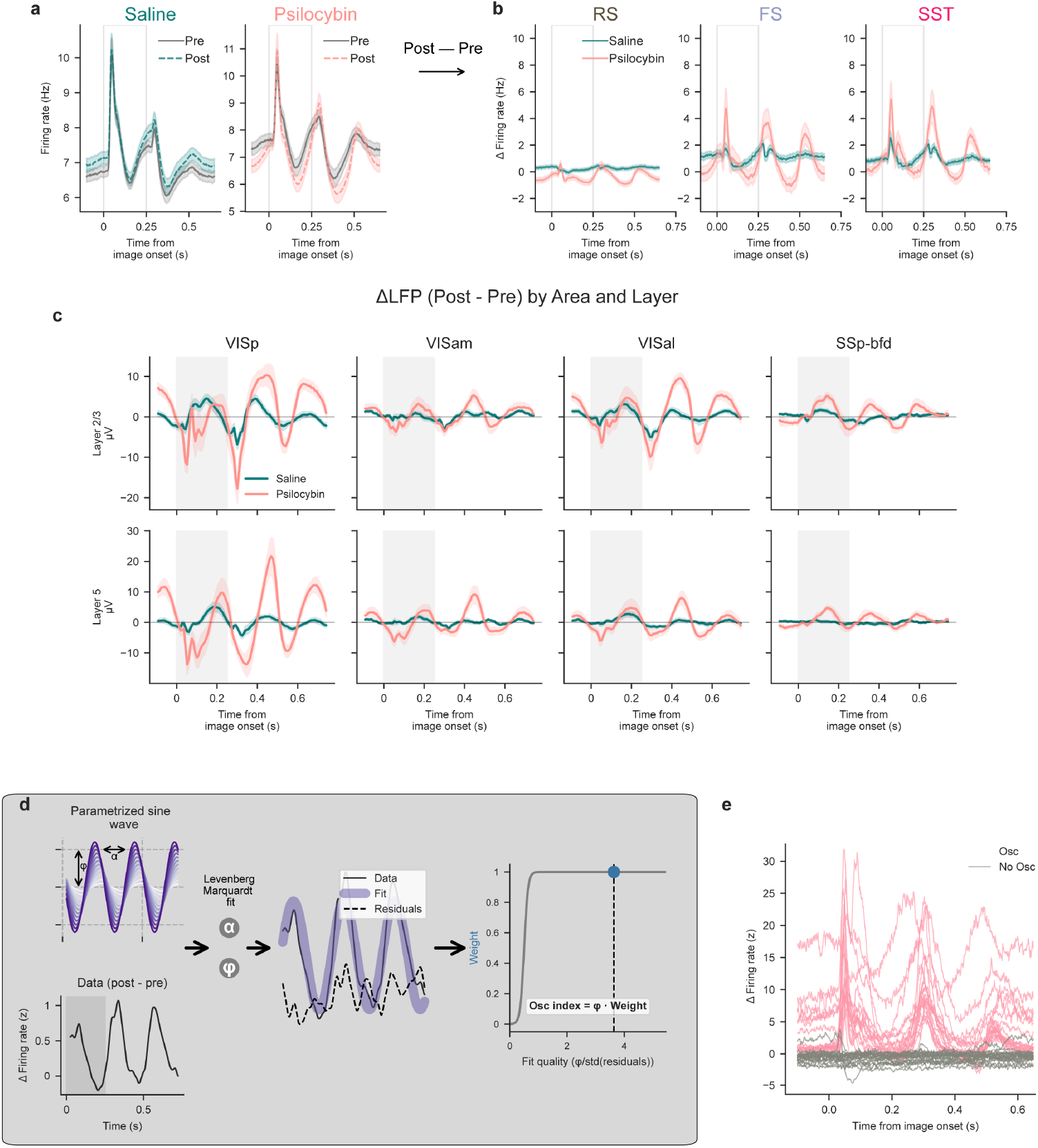
Psilocybin induces a a 4-Hz oscillation in visual cortex across scales. (a) Average PSTH of all cortical neurons aligned to all non-change images pre and post injection. Psilocybin causes a rhythmic change in firing rate. Shading indicates *±* 1 s.e.m. (b) ΔPSTH (post - pre epochs) of all cortical neurons recorded separately for each cell type. A clear modulation is visible across cell types, prominently in FS and SST neurons. Shading indicates *±* 1 s.e.m. (c) Layer 2/3 (top row) and layer 5 (bottom row) delta LFPs of VISp, VISam, VISal and SSp-bfd. Each delta LFP is computed by subtracting the image onset-aligned (only non-change)traces of the pre-injection block from the post-injection block. The psilocybin group shows a clear modulation similar to the one shown in the spike analysis. Shading indicates *±* 1 s.e.m. (d) Analytical framework for calculating the oscillation index. A 4-Hz sine wave was fitted to the delta PSTH (post-injection minus pre-injection epochs) of each individual neuron by optimizing both amplitude and phase. The resulting residuals were used to calculate a fit-quality metric, defined as the ratio of the fitted amplitude to the standard deviation of the residuals. To ensure the index selectively reflects robust rhythmic activity, the final oscillation index was down-weighted in cases of low fit quality (see Methods). (e) Example neurons from session ecephys_752311_2025-01-23_14-00-04. Osc index can reliably identify strongly oscillating neurons.

**Fig. S5.**
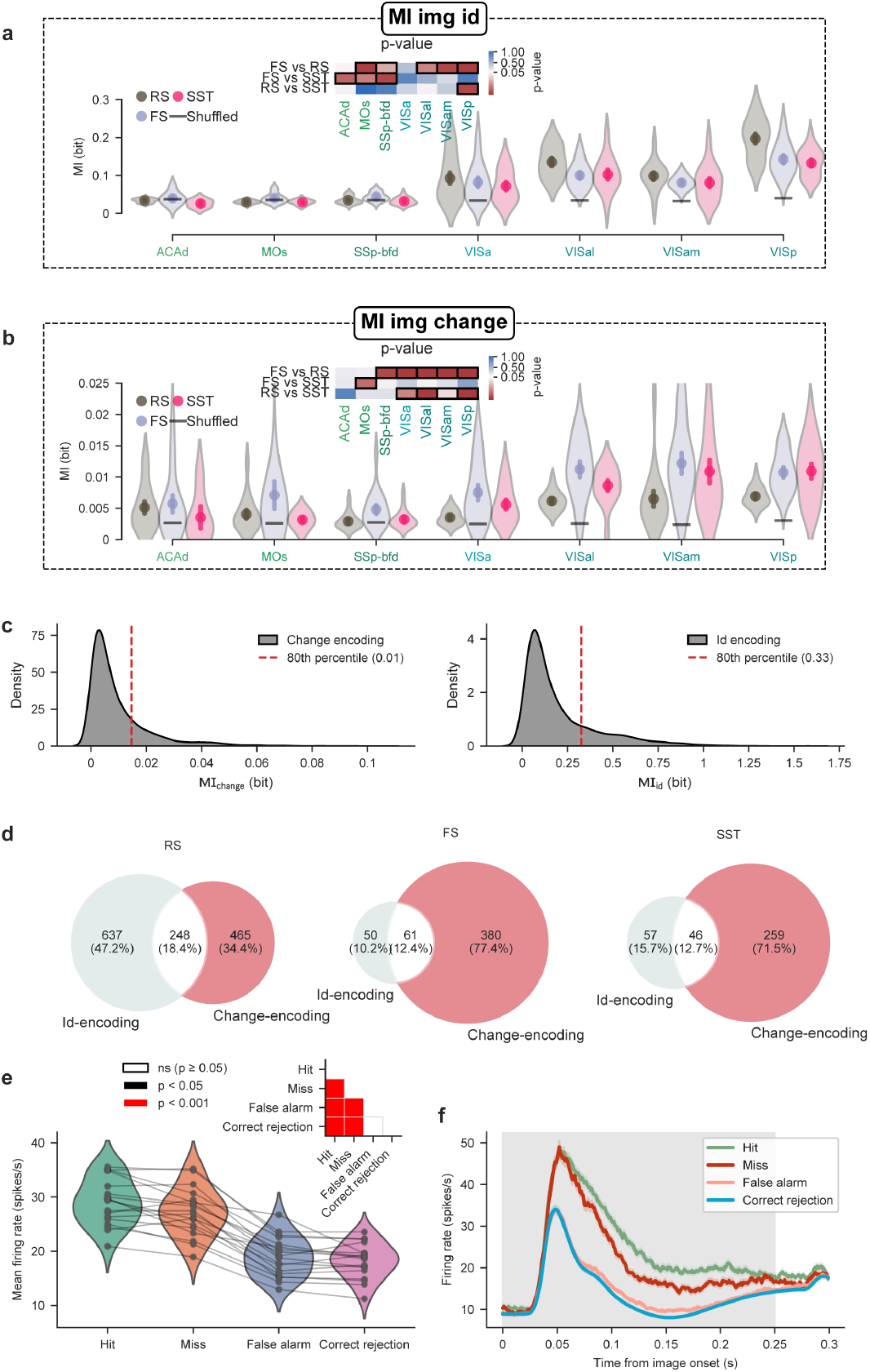
Encoding of image id and change across cortical neurons. (a) MI_id_ enrichment across visual areas and neuron types. Inset matrix shows p-values for each comparison; red indicates p <0.05, with darker shades representing stronger significance, while blue-to-white transitions denote values approaching significance (hierarchical bootstrap accounting for session id). (b) MI_change_ enrichment in FS and SST neurons is significant across visual areas. Inset matrix shows p-values for each comparison; red indicates p <0.05, with darker shades representing stronger significance, while blue-to-white transitions denote values approaching significance (hierarchical bootstrap accounting for session id). (c) Distribution of MI_change_ (left) and MI_id_ (right) values across neuron types. (d) Venn diagrams showing the overlap between change- and id-encoding neurons across neuron types. (e) Mean firing rate of change-encoding neurons across trial types (correct rejection = 10.8 *±* 0.3 spikes/s versus false alarm = 11.3 *±* 0.3 spikes/s versus hit = 14.6 *±* 0.4 spikes/s versus miss = 13.7 *±* 0.5 spikes/s; mean *±* s.e.m.; p-values determined by pairwise Wilcoxon signed-rank tests with Bonferroni correction). Inset shows the p-value significance matrix. (f) PSTHs of change-encoding neurons stratified by trial type.

**Fig. S6.**
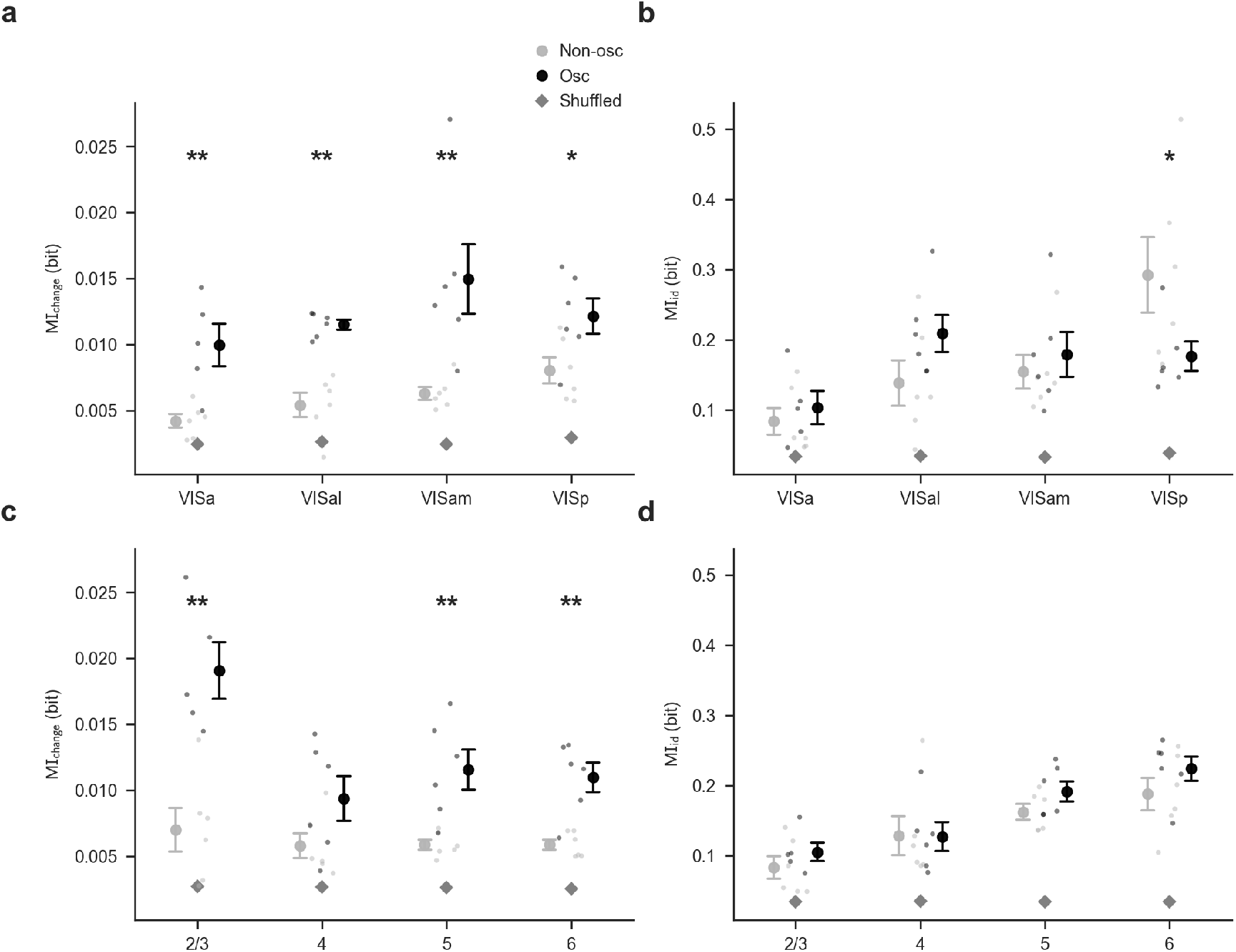
Change information is preferentially enriched in psilocybin-modulated cells across visual areas and cortical layers. (a) MI_change_ is enriched in psilocybin-modulated cells across brain areas (VISa, p=0.039, VISal, p<0.001, VISam, p<0.001, VISp, p=0.016, hierarchical bootstrap). Psilocybin-modulated neurons are defined as neurons with an Osc index in the top 20% percentile. (b) No widespread differences in MI_id_ by brain area, with the exception of VISp showing significantly reduced MI_id_ in psilocybin-modulated cells (VISa, p=0.93, VISal, p=0.54, VISam, p=0.30, VISp, p=0.017, hierarchical bootstrap).(c) Bottom: MI_change_ is enriched in psilocybin-modulated neurons across cortical layers, with layer 2/3 being particularly enriched, while layer 4 showing no enrichment (2/3, p<0.001; 4, p=0.14 5, p<0.001; 6, p<0.001, hierarchical bootstrap).(d) Bottom: MI_id_ across cortical layers, with layer 5 showing an enrichment in Osc neurons (2/3, p=0.37 4, p=0.99; 5, p<0.001; 6, p=0.45, hierarchical bootstrap).

**Fig. S7.**
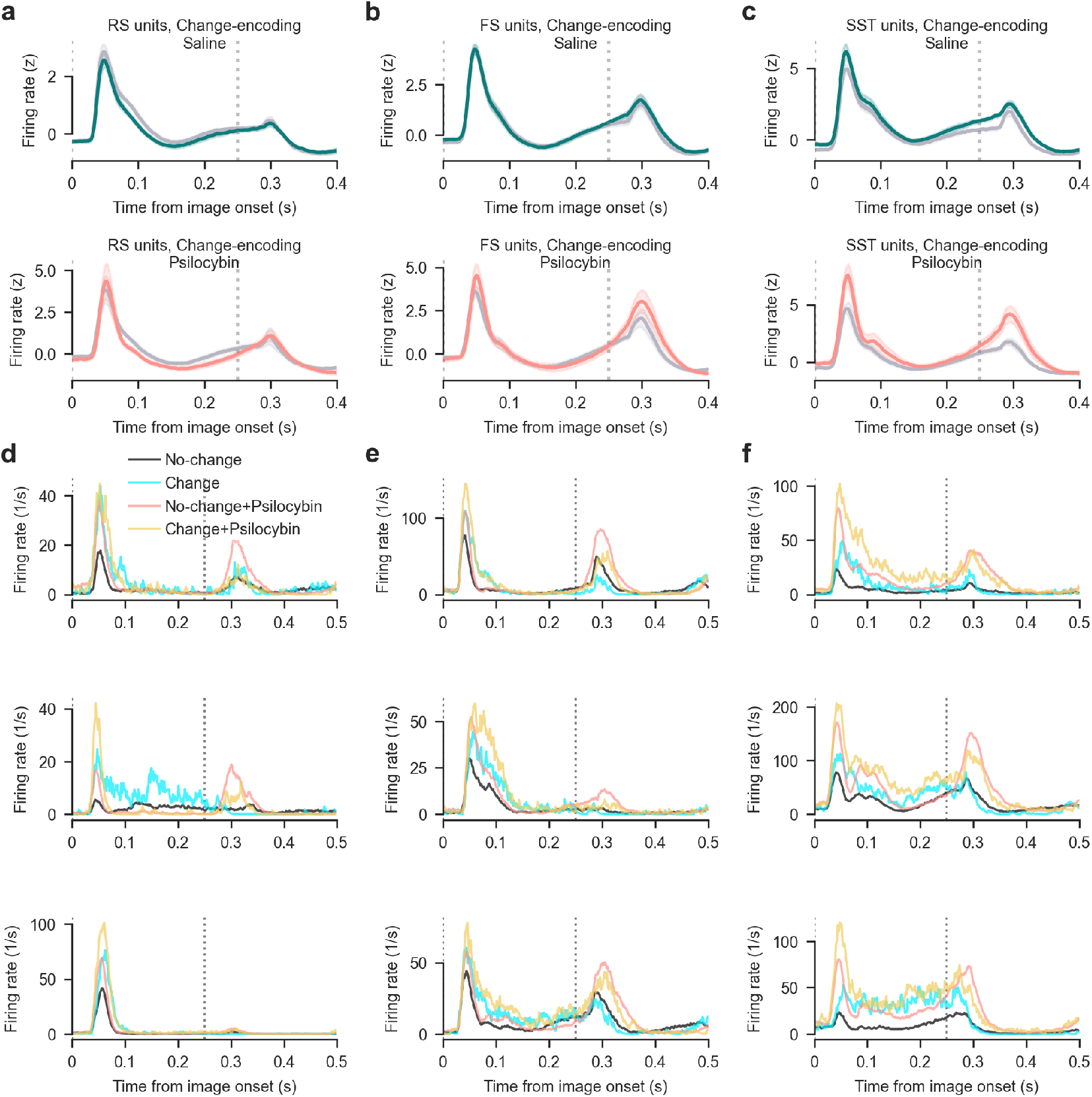
Change-encoding neurons’ responses across experimental conditions. (a) Top: responses of RS change-encoding neurons to no-change images before and after saline injection. Bottom: responses of RS change-encoding neurons to no-change images before and after psilocybin injection. (b) Same as (a) for FS change-encoding neurons. (c) Same as (a) for SST change-encoding neurons. (d) Responses of three single RS neurons to no-change (both before and after psilocybin injection) and change images. (d) Responses of three single FS neurons to no-change (both before and after psilocybin injection) and change images. (d) Responses of three single SST neurons to no-change (both before and after psilocybin injection) and change images.

**Fig. S8.**
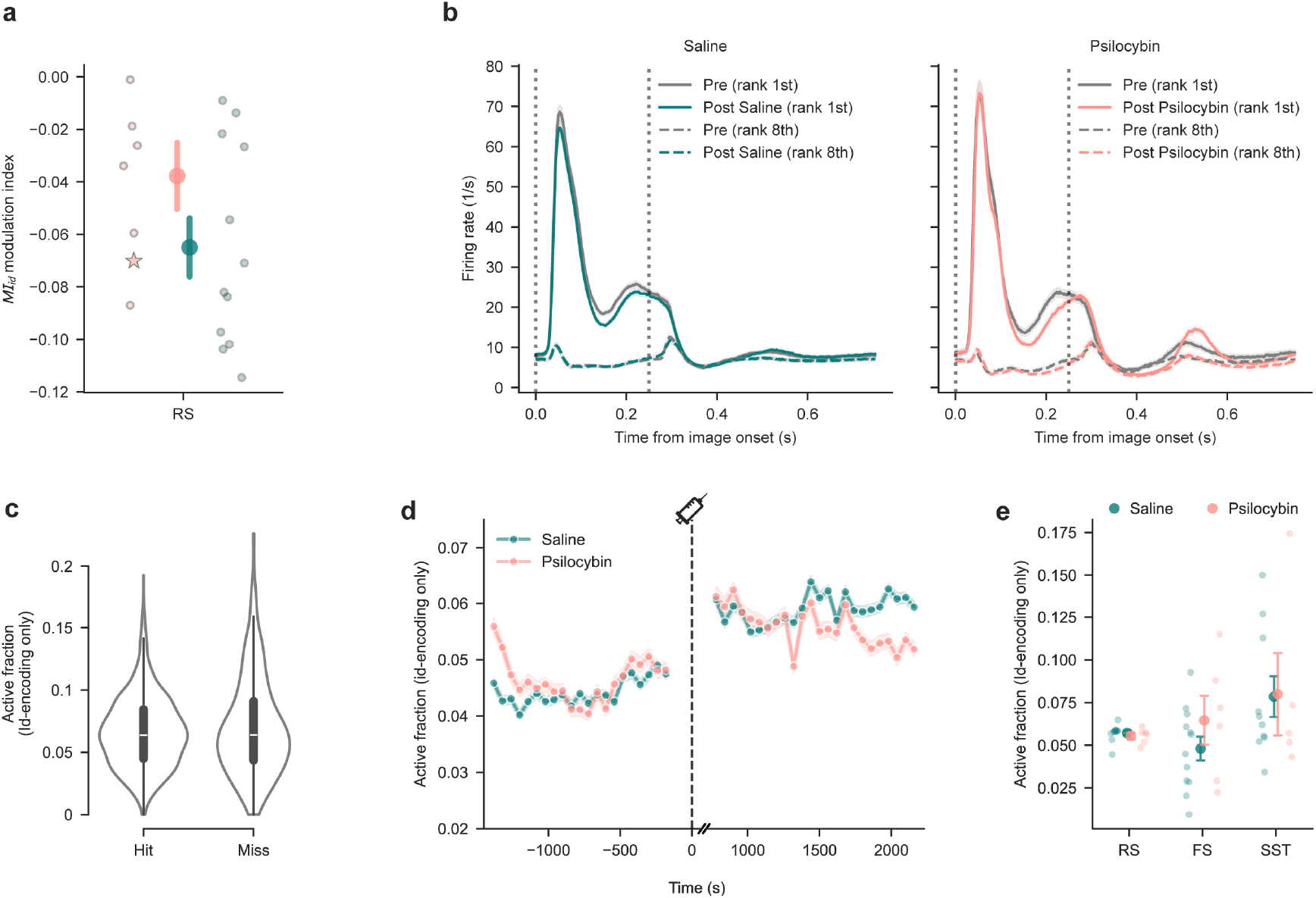
Image id-encoding neurons are not affected by psilocybin. (a) Average MI_id_ in image id-encoding neurons after injection, showing no significant change between saline and psilocybin conditions. (b) Population PSTHs of image id-encoding neurons to their preferred (rank 1) and least preferred (rank 8) image before and after injection of saline (left) or psilocybin (right). (c) Active fraction of image id-encoding neurons during change trials is not different between hit and miss trials. (d) Active fraction of image id-encoding neurons across before and after saline or psilocybin injection, showing no effect of psilocybin. (e) Active fraction is not different between groups in all three neuron types.

**Fig. S9.**
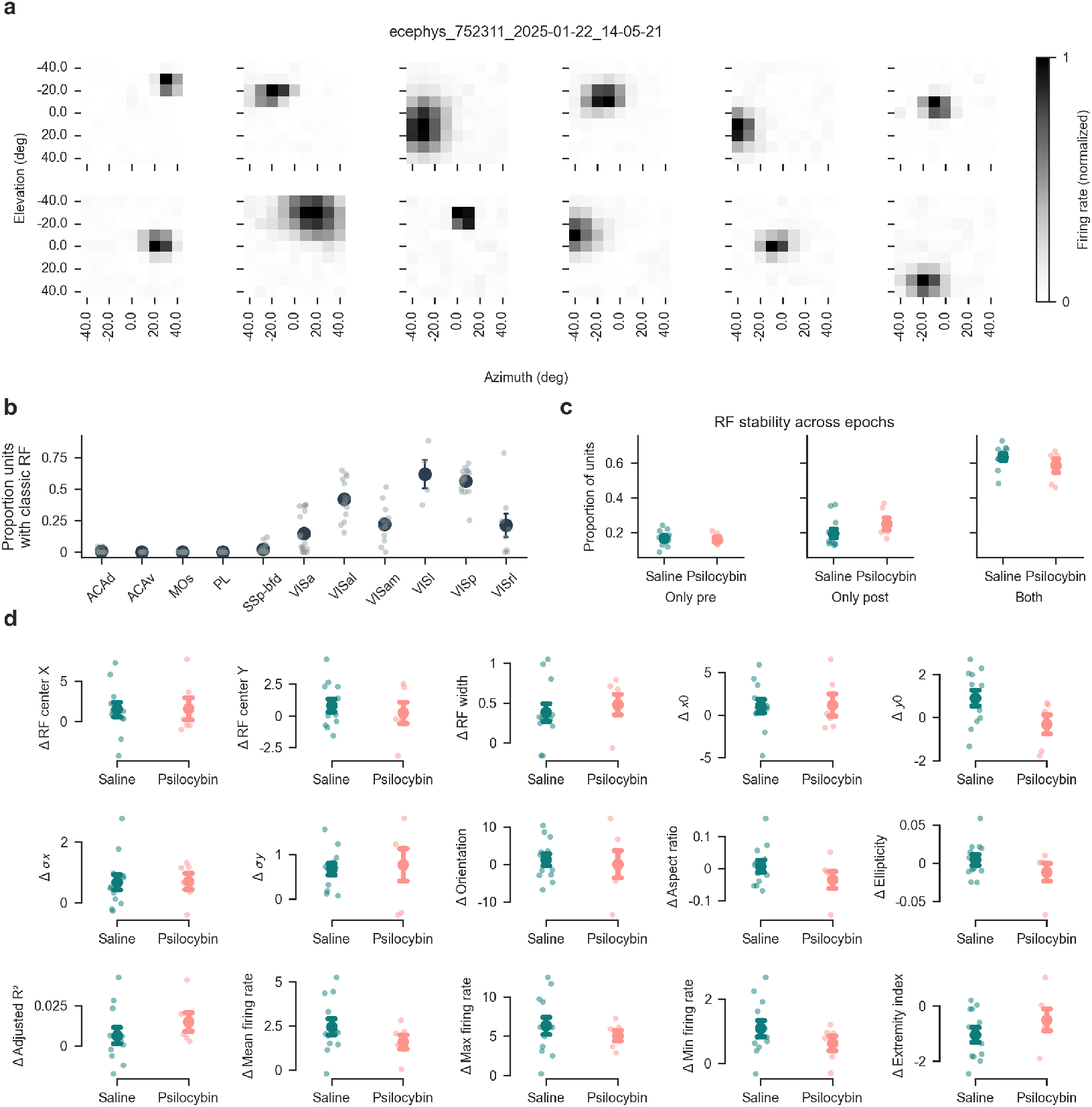
Receptive field tuning is not affected by psilocybin. (a) Representative examples of neurons with classic receptive field (RF) tuning. (b) Proportion of neurons exhibiting classical RF responses per brain area. (c) Proportion of neurons showing RF tuning only before injection (pre), only after injection (post) or in both epochs. Psilocybin does not affect the RF stability. (d) Psilocybin–evoked changes in RF parameters, compared to saline. Each panel shows the per-session mean change (*δ* = post - pre) for a given spatial, geometric, or baseline response metric. Extracted features encompass empirical properties (RF center x (medio-lateral), RF center y (dorso-ventral), and RF width), parametric variables optimized via rotated 2d Gaussian surface fits (x_0_, y_0_, *σ* x, *σ* y, orientation, aspect ratio, ellipticity, and adjusted R^2^), and global non-parametric matrix dynamics (extremity index, mean firing rate, min firing rate, and max firing rate). Each individual dot represents a single session average, and the point plots display the group mean *±* s.e.m. No significant differences in parameter dynamics were observed between the psilocybin and saline experimental cohorts (hierarchical bootstrap, all uncorrected p >0.05).

**Fig. S10.**
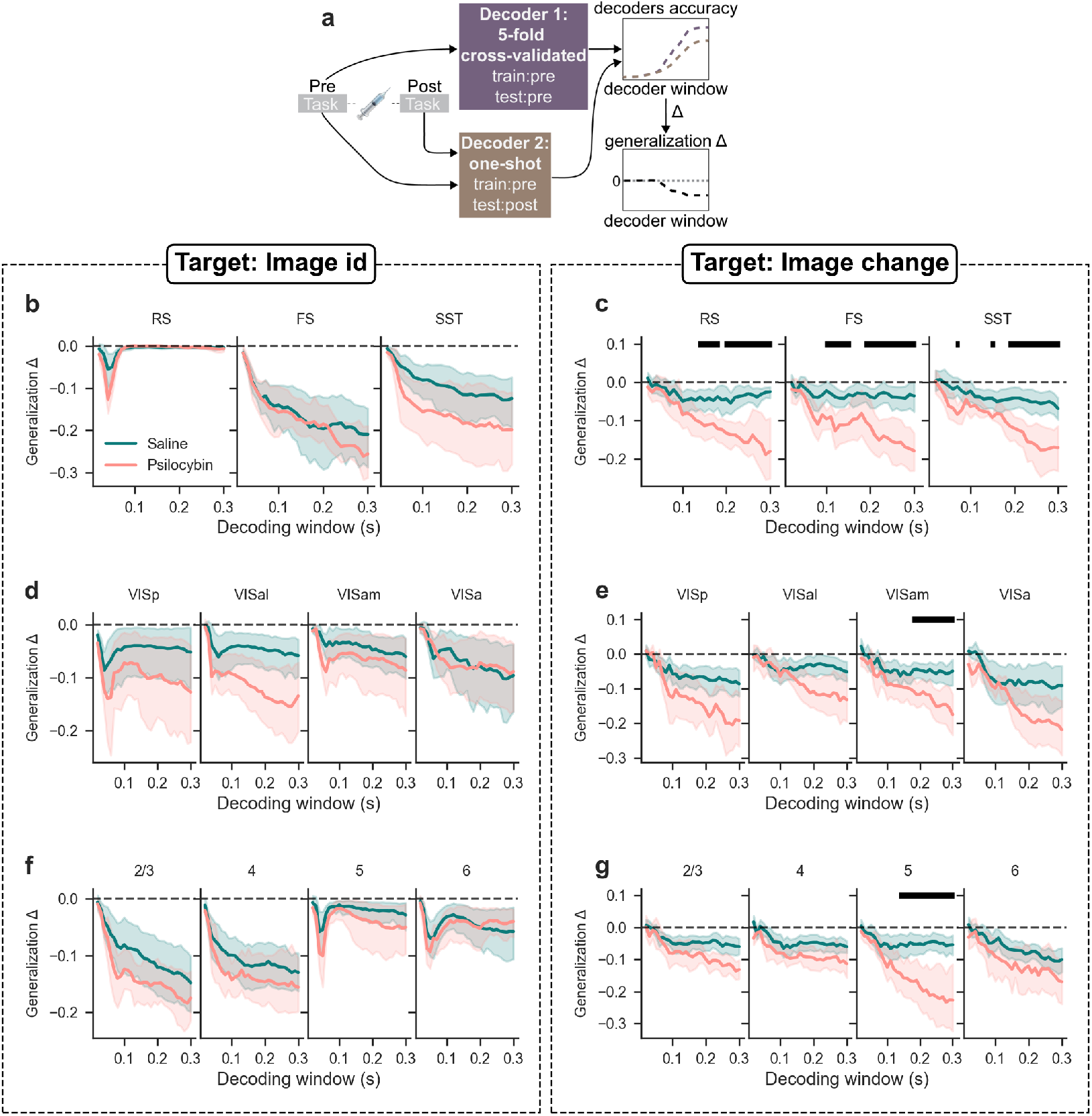
Decoding image identity and change across neuron types, brain areas and cortical layers. (a) Decoding procedure schematic. Two decoders are trained on activity during pre-injection epoch. The first one is tested using cross-validation on the same pre epoch, the second one is tested on the post epoch. (b) Decoding accuracy for image id over increasingly large time windows across neuron types. Black horizontal bars indicate significant differences between injection conditions (p <0.05, FDR-corrected independent two-sample t-tests). (c) Decoding accuracy for image change over increasingly large time windows across neuron types. (d) Decoding accuracy for image id over increasingly large time windows across brain areas. (e) Decoding accuracy for image change over increasingly large time windows across brain areas. (f) Decoding accuracy for image id over increasingly large time windows across cortical layers. (g) Decoding accuracy for image change over increasingly large time windows across cortical layers.

**Fig. S11.**
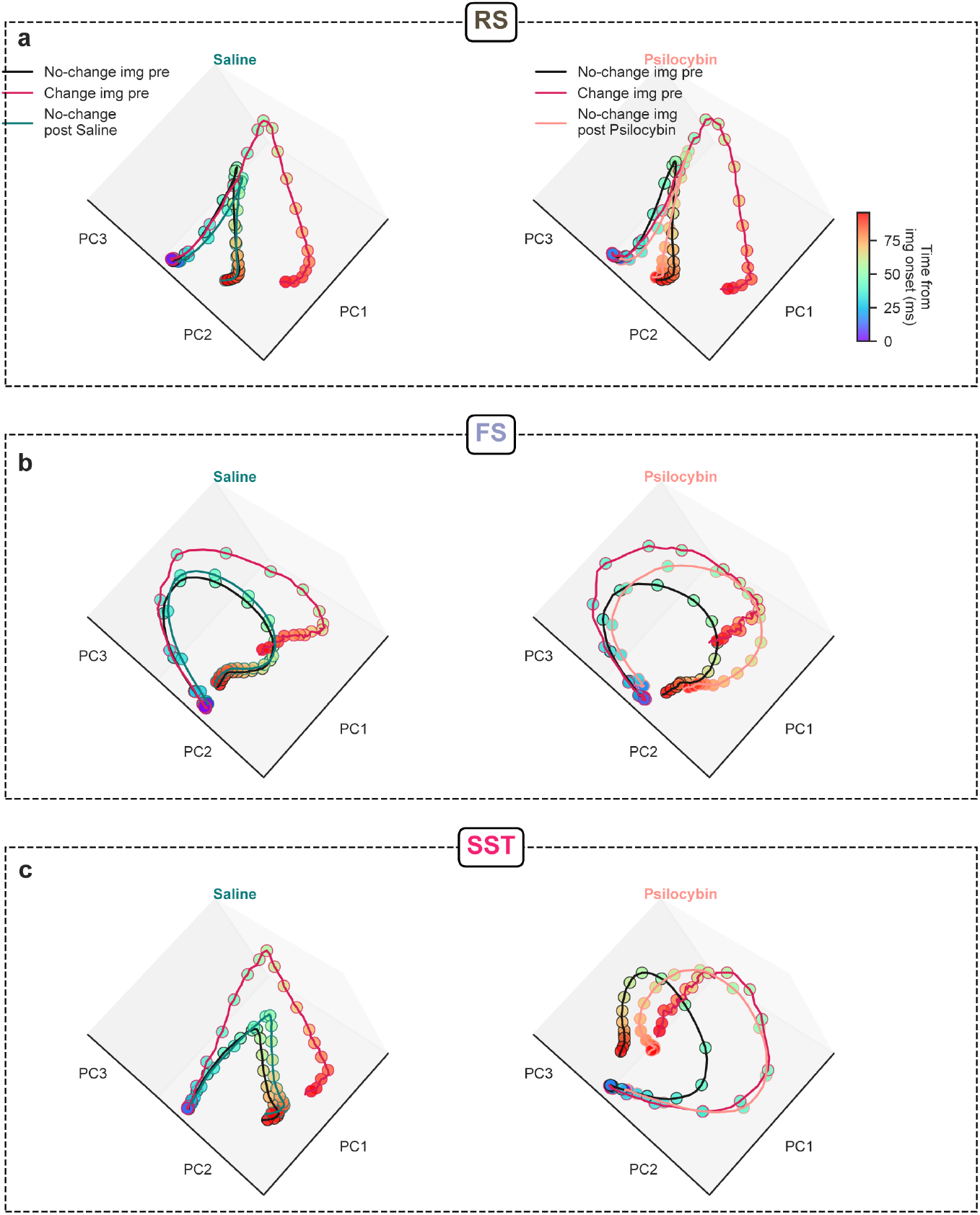
Psilocybin shifts sensory representation of non-change images in SST neurons. (a) RS-neuron population trajectory in the space defined by the first three principal components (PCs), showing RS activity during no-change images (black line), change images (orange line) and no-change images after injection (saline in teal on the left column, psilocybin in salmon on right column). Superimposed dots represent time progress from image onset 0-100 ms (white to black). (b) Same as (a) for FS neurons. (c) Same as (a) for SST neurons.

## Acknowledgments.

This work was supported by the European Research Council (ERC) under the European Union’s Horizon Europe research and innovation programme [grant agreement number 101117587, TIMEVALUE (TO)]. The experimental dataset in this project was obtained as part of the OpenScope program, which is operated by the Allen Institute, Neural Dynamics program and funded by the US National Institutes of Health (NIH) [grant numbers: U24NS113646 (JAL, CK)]. We thank the OpenScope steering committee and Karel Svoboda for their support. This research was also supported by the Allen Institute, founded by Jody Allen – chair and cofounder of Allen Family Philanthropies, and the late Paul G. Allen – investor, philanthropist, and co-founder of Microsoft. We gratefully acknowledge their vision and generosity, which make this work possible. We thank members of Ott lab, Dennis Nestvogel and Loreen Hertäg for discussion and Dietmar Schmitz for supporting the project.

## Declarations

### Author contributions

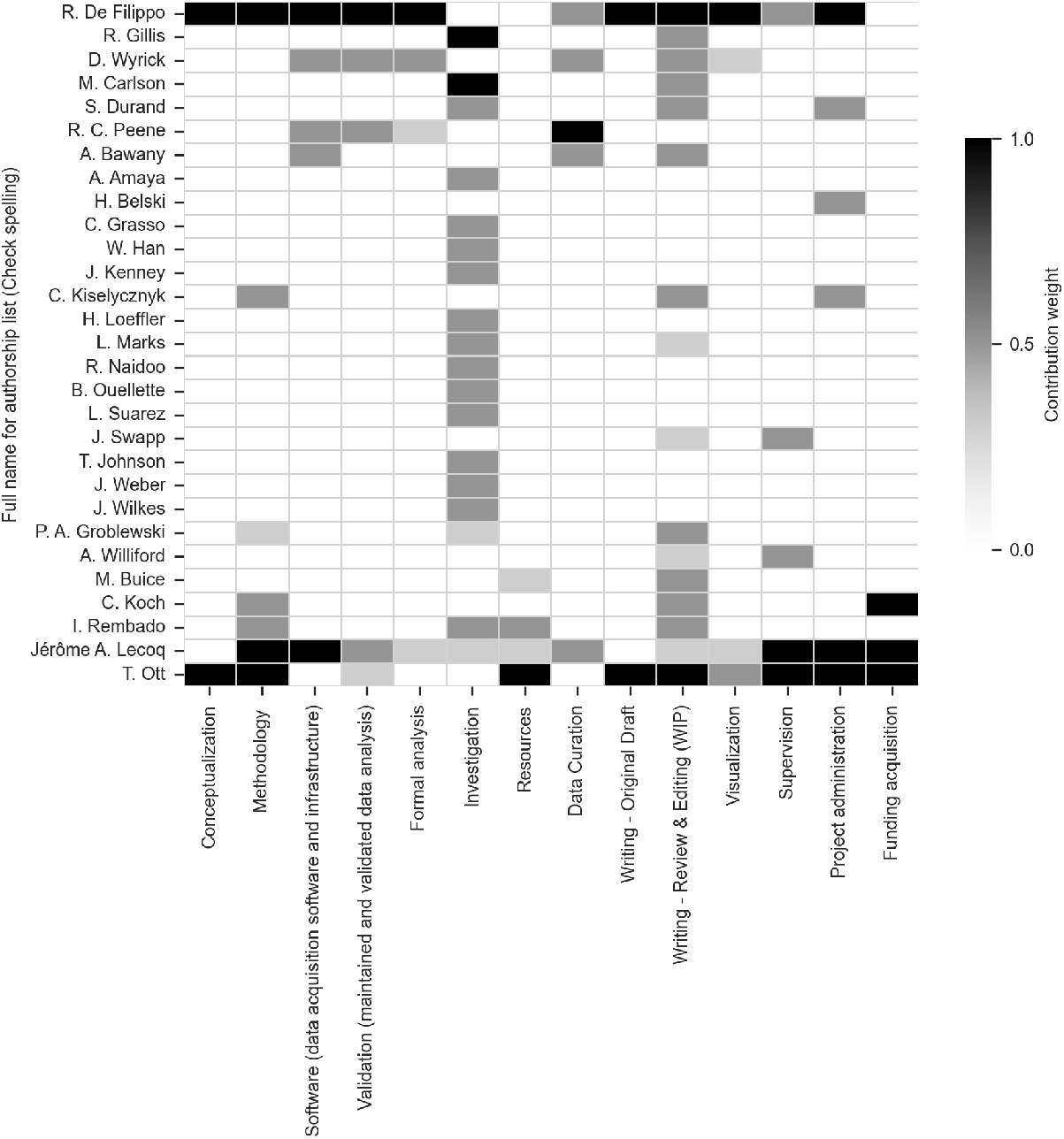

### Availability of data and materials

All the code used to process the dataset is available at https://github.com/RobertoDF/psycode. All figures and text can be reproduced using code present in this repository. Access to the original datasets is provided on the Dandi platform https://dandiarchive.org/dandiset/001417.

## References

[1] Key, B. J. Effects of Chlorpromazine and Lysergic Acid Diethylamide on the Rate of Habituation of the Arousal Response. Nature 190, 275–277 (1961). URL 10.1038/190275a0.

[2] Terrill, J. The nature of the LSD experience. The Journal of nervous and mental disease 135, 425–9 (1962). URL https://journals.lww.com/jonmd/Citation/1962/11000/The_Nature_of_the_LSD_Experience.6.aspx.

[3] Kometer, M. & Vollenweider, F. X. Serotonergic Hallucinogen-Induced Visual Perceptual Alterations, 257–282 (Springer Berlin Heidelberg, Berlin, Heidelberg, 2018). URL 10.1007/7854_2016_461.

[4] Carhart-Harris, R. L. & Friston, K. J. Rebus and the Anarchic Brain: Toward a Unified Model of the Brain Action of Psychedelics. Pharmacol Rev 71, 316–344 (2019). URL 10.1124/pr.118.017160.

[5] Corlett, P. R. et al. Hallucinations and Strong Priors. Trends in Cognitive Sciences 23, 114–127 (2019). URL 10.1016/j.tics.2018.12.001.

[6] De Filippo, R. & Schmitz, D. Synthetic Surprise as the Foundation of the Psychedelic Experience. Neuroscience & Biobehavioral Reviews 105538 (2024). URL 10.1016/j.neubiorev.2024.105538.

[7] Siegle, J. H. et al. Survey of spiking in the mouse visual system reveals functional hierarchy. Nature 592, 86–92 (2021). URL 10.1038/s41586-020-03171-x.

[8] Groblewski, P. A. et al. Characterization of Learning, Motivation, and Visual Perception in Five Transgenic Mouse Lines Expressing GCaMP in Distinct Cell Populations. Frontiers in Behavioral Neuroscience **Volume** 14 - 2020 (2020). URL 10.3389/fnbeh.2020.00104.

[9] Attinger, A., Wang, B. & Keller, G. B. Visuomotor coupling shapes the functional development of mouse visual cortex. Cell 169, 1291–1302. e14 (2017). URL 10.1016/j.cell.2017.05.023.

[10] Green, J. et al. A cell-type-specific error-correction signal in the posterior parietal cortex. Nature 620, 366–373 (2023). URL 10.1038/s41586-023-06357-1.

[11] Keller, A. J., Roth, M. M. & Scanziani, M. Feedback generates a second receptive field in neurons of the visual cortex. Nature 582, 545–549 (2020). URL 10.1038/s41586-020-2319-4.

[12] Adesnik, H., Bruns, W., Taniguchi, H., Huang, Z. J. & Scanziani, M. A neural circuit for spatial summation in visual cortex. Nature 490, 226–231 (2012). URL 10.1038/nature11526.

[13] Bennett, C. et al. Map of spiking activity underlying change detection in the mouse visual system. *bioRxiv* 2025.10.17.683190 (2025). URL 10.1101/2025.10.17.683190.

[14] Claar, L. D. et al. Cortico-thalamo-cortical interactions modulate electrically evoked EEG responses in mice. eLife 12, RP84630 (2023). URL 10.7554/eLife.84630.

[15] Holze, F. et al. Direct comparison of the acute effects of lysergic acid diethylamide and psilocybin in a double-blind placebo-controlled study in healthy subjects. Neuropsychopharmacology 47, 1180–1187 (2022). URL 10.1038/s41386-022-01297-2.

[16] Weinreb, C. et al. Keypoint-MoSeq: parsing behavior by linking point tracking to pose dynamics. Nature Methods 21, 1329–1339 (2024). URL 10.1038/s41592-024-02318-2.

[17] Purple, R. J. et al. Short-and long-term modulation of rat prefrontal cortical activity following single doses of psilocybin. Molecular Psychiatry 30, 5889–5900 (2025). URL 10.1038/s41380-025-03182-y.

[18] Skyberg, R. J., Fields, C. W., Martins, D. M. & Niell, C. M. The impact of the serotonergic psychedelic DOI on active vision in freely moving mice. *bioRxiv* (2025). URL 10.1101/2025.10.14.682230.

[19] Michaiel, A. M., Parker, P. R. L. & Niell, C. M. A Hallucinogenic Serotonin-2a Receptor Agonist Reduces Visual Response Gain and Alters Temporal Dynamics in Mouse V1. Cell Rep 26, 3475–3483.e4 (2019). URL 10.1016/j.celrep.2019.02.104.

[20] Syeda, A. et al. Facemap: a framework for modeling neural activity based on orofacial tracking. Nature Neuroscience 27, 187–195 (2024). URL 10.1038/s41593-023-01490-6.

[21] Bennett, C. et al. Shield: Skull-shaped hemispheric implants enabling large-scale electrophysiology datasets in the mouse brain. Neuron 112, 2869–2885.e8 (2024). URL 10.1016/j.neuron.2024.06.015.

[22] Harris, J. A. et al. Hierarchical organization of cortical and thalamic connectivity. Nature 575, 195–202 (2019). URL 10.1038/s41586-019-1716-z.

[23] Stringer, C., Michaelos, M., Tsyboulski, D., Lindo, S. E. & Pachitariu, M. High-precision coding in visual cortex. Cell 184, 2767–2778.e15 (2021). URL 10.1016/j.cell.2021.03.042.

[24] White, C. M. et al. Psychedelic 5-HT2A agonist increases spontaneous and evoked 5-Hz oscillations in visual and retrosplenial cortex. Communications Biology 9, 216 (2026). URL 10.1038/s42003-025-09492-9.

[25] Nestvogel, D. B. & McCormick, D. A. Visual thalamocortical mechanisms of waking state-dependent activity and alpha oscillations. Neuron 110, 120–138.e4 (2022). URL 10.1016/j.neuron.2021.10.005.

[26] Momi, D. et al. Brain-wide reconfiguration of burst firing by psilocybin reveals 5-HT2A-dependent circuit dynamics. *bioRxiv* 2026.08.14.744865 (2026). URL 10.64898/2026.08.14.744865.

[27] Shao, L. X. et al. Psilocybin induces rapid and persistent growth of dendritic spines in frontal cortex in vivo. Neuron 109, 2535–2544.e4 (2021). URL 10.1016/j.neuron.2021.06.008.

[28] Brockett, A. T. & Francis, N. A. Psilocybin decreases neural responsiveness and increases functional connectivity while preserving pure-tone frequency selectivity in mouse auditory cortex. J Neurophysiol 132, 45–53 (2024). URL 10.1152/jn.00124.2024.

[29] Halberstadt, A. L., Chatha, M., Klein, A. K., Wallach, J. & Brandt, S. D. Correlation between the potency of hallucinogens in the mouse head-twitch response assay and their behavioral and subjective effects in other species. Neuropharmacology 167, 107933 (2020). URL 10.1016/j.neuropharm.2019.107933.

[30] Bradley, P. B., Elkes, C. & Elkes, J. On some effects of lysergic acid diethylamide (L.S.D. 25) in normal volunteers. J Physiol 121, 50p–51p (1953). URL https://europepmc.org/article/MED/13085360.

[31] Roberts, B. F. et al. Effect of psilocybin on decision-making and motivation in the healthy rat. Behav Brain Res 440, 114262 (2023). URL 10.1016/j.bbr.2022.114262.

[32] Torrado Pacheco, A., Olson, R. J., Garza, G. & Moghaddam, B. Acute psilocybin enhances cognitive flexibility in rats. Neuropsychopharmacology 48, 1011–1020 (2023). URL 10.1038/s41386-023-01545-z.

[33] Schmitz, G. P. et al. Psychedelic compounds directly excite 5-HT2A layer V medial prefrontal cortex neurons through 5-HT2A Gq activation. Translational Psychiatry 15, 381 (2025). URL 10.1038/s41398-025-03611-0.

[34] Ekins, T. G. et al. Cellular rules underlying psychedelic control of prefrontal pyramidal neurons. bioRxiv (2023). URL 10.1101/2023.10.20.563334.

[35] Vargas, M. V. et al. Psychedelics promote neuroplasticity through the activation of intracellular 5-HT2A receptors. Science 379, 700–706 (2023). URL 10.1126/science.adf0435.

[36] Furutachi, S., Franklin, A. D., Aldea, A. M., Mrsic-Flogel, T. D. & Hofer, S. B. Cooperative thalamocortical circuit mechanism for sensory prediction errors. Nature 633, 398–406 (2024). URL 10.1038/s41586-024-07851-w.

[37] Hamm, J. P. & Yuste, R. Somatostatin Interneurons Control a Key Component of Mismatch Negativity in Mouse Visual Cortex. Cell Reports 16, 597–604 (2016). URL 10.1016/j.celrep.2016.06.037.

[38] Bastos, G., et al. Top-down input modulates visual context processing through an interneuron-specific circuit. Cell Reports 42 (2023). URL 10.1016/j.celrep.2023.113133.

[39] Natan, R. G. et al. Complementary control of sensory adaptation by two types of cortical interneurons. eLife 4, e09868 (2015). URL 10.7554/eLife.09868.

[40] Iigaya, K., Fonseca, M. S., Murakami, M., Mainen, Z. F. & Dayan, P. An effect of serotonergic stimulation on learning rates for rewards apparent after long intertrial intervals. Nat Commun 9, 2477 (2018). URL 10.1038/s41467-018-04840-2.

[41] Matias, S., Lottem, E., Dugué, G. P. & Mainen, Z. F. Activity patterns of serotonin neurons underlying cognitive flexibility. eLife 6, e20552 (2017). URL 10.7554/eLife.20552.

[42] Grossman, C. D., Bari, B. A. & Cohen, J. Y. Serotonin neurons modulate learning rate through uncertainty. Curr Biol 32, 586–599.e7 (2022). URL 10.1016/j.cub.2021.12.006.

[43] Hubert, F. et al. A state prediction error model explains serotonin activity and its role in cognitive flexibility. Cosyne Abstracts 2026 (2026).

[44] Zhang, M. et al. Molecularly defined and spatially resolved cell atlas of the whole mouse brain. Nature 624, 343–354 (2023). URL 10.1038/s41586-023-06808-9.

[45] Yao, Z. et al. A high-resolution transcriptomic and spatial atlas of cell types in the whole mouse brain. Nature 624, 317–332 (2023). URL 10.1038/s41586-023-06812-z.

[46] De Filippo, R. & Schmitz, D. Transcriptomic mapping of the 5-HT receptor landscape. Patterns URL 10.1016/j.patter.2024.101048.

[47] Davoudian, P. A. et al. Psilocybin reshapes cortical inhibition through selective interneuron recruitment. bioRxiv 2026.04.16.718963 (2026). URL 10.64898/2026.04.16.718963.

[48] Horrocks, M., Mohn, J. L. & Jaramillo, S. The serotonergic psychedelic DOI impairs deviance detection in the auditory cortex. J Neurophysiol 133, 388–398 (2025). URL 10.1152/jn.00411.2024.

[49] Timmermann, C. et al. Lsd modulates effective connectivity and neural adaptation mechanisms in an auditory oddball paradigm. Neuropharmacology 142, 251–262 (2018). URL 10.1016/j.neuropharm.2017.10.039.

[50] Duerler, P. et al. Psilocybin Induces Aberrant Prediction Error Processing of Tactile Mismatch Responses—A Simultaneous EEG–FMRI Study. Cerebral Cortex 32, 186–196 (2022). URL 10.1093/cercor/bhab202.

[51] Martin, D. A., Delgado, A. M. & Calu, D. J. Effects of psychedelic, DOI, on nucleus accumbens dopamine signaling to predictable rewards and cues in rats. Neuropsychopharmacology 49, 1925–1933 (2024). URL 10.1038/s41386-024-01912-4.

[52] Key, B. J. & Bradley, P. B. Effect of Drugs on Conditioning and Habituation to Arousal Stimuli in Animals. Nature 182, 1517–1519 (1958). URL 10.1038/1821517a0.

[53] Lu, O. D. et al. A multi-institutional investigation of psilocybin’s effects on mouse behavior. bioRxiv 2025.04.08.647810 (2025). URL 10.1101/2025.04.08.647810.

[54] Barrett, L. F., Quigley, K. S. & Hamilton, P. An active inference theory of allostasis and interoception in depression. Philosophical Transactions of the Royal Society B: Biological Sciences 371, 20160011 (2016). URL 10.1098/rstb.2016.0011.

[55] Kube, T., Schwarting, R., Rozenkrantz, L., Glombiewski, J. A. & Rief, W. Distorted Cognitive Processes in Major Depression: A Predictive Processing Perspective. Biological Psychiatry 87, 388–398 (2020). URL 10.1016/j.biopsych.2019.07.017.

[56] Strube, A. & Pizzagalli, D. A. Brain rhythms of depression: A predictive processing perspective. Trends Neurosci (2026). URL 10.1016/j.tins.2026.06.006.

[57] Fee, C., Banasr, M. & Sibille, E. Somatostatin-Positive Gamma-Aminobutyric Acid Interneuron Deficits in Depression: Cortical Microcircuit and Therapeutic Perspectives. Biol Psychiatry 82, 549–559 (2017). URL 10.1016/j.biopsych.2017.05.024.

[58] Guo, H. et al. Somatostatin-expressing neurons in the zona incerta regulate chronic stress response and modulate depression-like behaviors. Molecular Psychiatry (2026). URL 10.1038/s41380-026-03446-1.

[59] Fuchs, T. et al. Disinhibition of somatostatin-positive GABAergic interneurons results in an anxiolytic and antidepressant-like brain state. Mol Psychiatry 22, 920–930 (2017). URL 10.1038/mp.2016.188.

[60] Anderson, K. M. et al. Convergent molecular, cellular, and cortical neuroimaging signatures of major depressive disorder. Proceedings of the National Academy of Sciences 117, 25138–25149 (2020). URL 10.1073/pnas.2008004117.

[61] Prévot, T. & Sibille, E. Altered GABA-mediated information processing and cognitive dysfunctions in depression and other brain disorders. Molecular Psychiatry 26, 151–167 (2021). URL 10.1038/s41380-020-0727-3.

[62] West, C. L. et al. Psychedelics relax predictive processing in the post-acute period by remodeling cortico-cortical feedback circuits. *bioRxiv* 2024.07.03.601959 (2026). URL 10.1101/2024.07.03.601959.

[63] Shao, L. X. et al. Pyramidal cell types and 5-HT(2a) receptors are essential for psilocybin’s lasting drug action. bioRxiv (2024). URL 10.1101/2024.11.02.621692.

[64] Ly, C. et al. Psychedelics Promote Structural and Functional Neural Plasticity. Cell Reports 23, 3170–3182 (2018). URL 10.1016/j.celrep.2018.05.022.

[65] Buccino, A. P. et al. Spikeinterface, a unified framework for spike sorting. Elife 9 (2020). URL 10.7554/eLife.61834.

[66] Birman, D. et al. Pinpoint: trajectory planning for multi-probe electrophysiology and injections in an interactive web-based 3d environment (2023). URL 10.7554/elife.91662.1.

[67] Juavinett, A. L., Nauhaus, I., Garrett, M. E., Zhuang, J. & Callaway, E. M. Automated identification of mouse visual areas with intrinsic signal imaging. Nat Protoc 12, 32–43 (2017). URL 10.1038/nprot.2016.158.

[68] Durand, S. et al. Acute head-fixed recordings in awake mice with multiple Neuropixels probes. Nat Protoc 18, 424–457 (2023). URL 10.1038/s41596-022-00768-6.

[69] Jun, J. J. et al. Fully integrated silicon probes for high-density recording of neural activity. Nature 551, 232–236 (2017). URL 10.1038/nature24636.

[70] Siegle, J. H. et al. Open Ephys: an open-source, plugin-based platform for multichannel electrophysiology. J Neural Eng 14, 045003 (2017). URL 10.1088/1741-2552/aa5eea.

[71] Pachitariu, M., Sridhar, S., Pennington, J. & Stringer, C. Spike sorting with Kilosort4. Nature Methods 21, 914–921 (2024). URL 10.1038/s41592-024-02232-7.

[72] Jain, A. et al. Unitrefine: A Community Toolbox for Automated Spike Sorting Curation. *bioRxiv* 2025.03.30.645770 (2025). URL 10.1101/2025.03.30.645770.

[73] Myers, H. & Toglia, D. Refractive Index Matching - EasyIndex v1 (2023). URL http://dx.doi.org/10.17504/protocols.io.kxygx965kg8j/v1.

[74] Mathis, A. et al. Deeplabcut: markerless pose estimation of user-defined body parts with deep learning. Nature Neuroscience 21, 1281–1289 (2018). URL 10.1038/s41593-018-0209-y.

[75] Pedregosa, F. et al. Scikit-learn: Machine learning in Python. the Journal of machine Learning research 12, 2825–2830 (2011). URL https://jmlr.org/papers/v12/pedregosa11a.html.

